# Systematic Spike Sorting Errors Revealed by Juxtacellular Ground Truth in the Awake Brain

**DOI:** 10.64898/2026.10.29.762918

**Authors:** Zach Saccomano, Behnaz Khaghani, Reagan Bullins Shugg, Sam McKenzie, Daniel F. English

**Author notes:** These authors contributed equally to this work (Zach Saccomano, Behnaz Khaghani). **Correspondence:** Daniel F. English.

## Abstract

Existing ground-truth data for spike sorting come almost entirely from anesthetized or in vitro preparations. Yet phenomena expected to challenge spike sorting most—spike overlap and bursting—depend on brain state and differ in the awake brain. Therefore, error estimates from existing benchmarks may not transfer to awake recordings, and the types of errors in the awake brain remain largely uncharacterized. To address this, we generated a ground-truth dataset (16 neurons) using paired juxtacellular-extracellular electrophysiology in awake mice, and found spike sorting performance, particularly recall, to be lower than conventional benchmarks might suggest. Generally, false negatives sharply increased with intrinsic physiological factors like bursting, whereas false positives were more strongly associated with extrinsic factors, such as putative spike overlaps. To quantify contributions to errors from intrinsic and extrinsic factors, for each observed juxtacellular waveform we predicted the corresponding extracellular waveform. In the best-performing sorter, we estimate approximately half of false positives were attributable to credulous attempts to resolve putative overlaps. Bursting and putative overlaps were each associated with a roughly doubled false negative rate when they occurred separately, and with a sevenfold increase when they occurred together. These co-occurrences accounted for approximately 40% of a typical neuron’s false negatives, despite comprising only approximately a fifth of its spikes. Thus, the moments when spike sorters must rely on the assumption of waveform invariance the most to resolve overlaps are exactly the moments when bursting makes that assumption least valid. We also estimate that three-quarters of false negatives had been detected but assigned to other units. These spikes were not cleanly oversplit: post-hoc merging was not productive because the loss in precision was three times the gain in recall. Because bursting and spike overlaps often co-occur, sorting spikes by waveform similarity and repairing the result by post hoc merging is unlikely to adequately balance precision and recall in the situations of greatest scientific interest. That is, while bursting and spike overlaps may involve only a minority of spikes, the resulting sorting errors may disproportionately distort the interpretation of neuronal population dynamics. We thus conclude that uncertainty quantification in sorters’ output is essential.

**Significance Statement:** Spike sorting, the assignment of extracellular spikes to individual neurons, underlies most large-scale recordings of brain activity, yet its accuracy has been validated almost entirely in simulations or in anesthetized or ex vivo preparations. Using simultaneous juxtacellular and extracellular recordings in awake mice, we show that conventional benchmarks likely overestimate sorting performance in the awake brain, and that sorting errors are highly structured. False negatives prominently concentrate where bursting coincides with spike overlap, and most appear to be detected but assigned to other, low-precision clusters, from which merging cannot cleanly recover them. Roughly half of false positives likely come from wrongly resolved spike overlaps. Because structured errors can masquerade as temporal relationships between neurons, they can disproportionately distort conclusions about neural coding, population dynamics, and functional connectivity. Our publicly available recordings provide a tool to devise more robust spike sorting assumptions, and our results argue for sorters that model burst-related waveform variability and report their uncertainty, especially during spike overlaps.

## Introduction

Quantifying and interpreting spike sorting errors requires ground truth: a recording in which the spike times of at least one neuron are known independently, and with greater confidence. Most existing biological ground-truth data have been obtained from ex vivo preparations (Anastassiou et al., 2015; Yger et al., 2018; Spampinato et al., 2018) or anesthetized animals (Harris et al., 2000; Henze et al., 2000, 2009; Neto et al., 2016; Allen et al., 2018; Marques-Smith et al., 2018), conditions in which neuronal activity may differ from activity in awake, drug-free animals. The rates and patterns of spike sorting errors observed under these conditions therefore need not generalize to awake recordings.

An alternative source of ground truth is simulation, either as hybrid data, in which curated templates are inserted at known times into real recordings, or as fully synthetic recordings generated from biophysical models (Martinez et al., 2009; Pachitariu et al., 2016; Rossant et al., 2016; Hagen et al., 2015; Wouters et al., 2021; Buccino and Einevoll, 2021). Simulations yield large sample sizes and permit controlled comparisons, but their relevance depends on how faithfully they reproduce biological variability, including burst-related wave-form variation and the temporal structure of population activity. Importantly, in practice, spike sorting simulations are often built on the sorters’ own assumptions, and are thus circular to that extent.

In contrast, today, perhaps the extracellular population recordings of greatest scientific interest are made in awake, behaving animals (Steinmetz et al., 2019; Stringer et al., 2019; International Brain Laboratory et al., 2024). This matters for benchmarking because neuronal dynamics and the statistical structure of population activity vary with brain state. Relevant phenomena include the biophysics of spiking, including burst-related changes in a neuron’s waveform (Steriade et al., 2001; Sherman, 2001; Steriade, 2004; Fee et al., 1996; Lewicki, 1998; Harris et al., 2000; Chung et al., 2017; Allen et al., 2018; Koukuntla et al., 2025); the population correlation structure, which changes how often spikes overlap (Ecker et al., 2014; Aasebø et al., 2017); and LFP state, which changes the background against which spikes are detected. Indeed, sorting success has been directly shown to differ between anesthetic states because of differences in coordinated population activity (Hildebrandt et al., 2017).

To characterize the failure modes of spike sorting in the awake brain, we recorded 16 neurons juxtacellularly in the neocortex, thalamus, CA1, and subiculum of awake mice with simultaneous 32-channel extracellular recording. Ground-truth labels are never exact: small-amplitude events within bursts can be continuous with the noise distribution rather than cleanly separable from it, and the awake preparation, being less stable, compounds this considerably. To account for this uncertainty, we assigned each candidate juxtacellular event a posterior probability of being a spike, and used the resulting distribution over labelings to compute soft performance metrics. Against this benchmark, we evaluated Kilosort4 (KS4), IronClust (IC), MountainSort5 (MS5), and SpyKING CIRCUS2 (SC2), and decomposed their errors by sources of variability intrinsic or extrinsic to the recorded neuron. We first focus on detailed analyses in the best-performing sorter (KS4) and then test key findings in the others. In a typical neuron, we found that 80% of burst-modulated spikes were recovered when no other neuron fired nearby, but burst-attenuated spikes with spike overlap-like residuals accounted for a disproportionate share of false negatives: a median of 40% of false negatives despite comprising 22% of spikes. We further estimate that, in a typical neuron, 75% of false negatives were detected but assigned to other units, while 45% of false positives appeared to arise from credulous attempts to resolve putative spike overlaps.

## Materials and Methods

Terms and abbreviations used throughout are collected in Table 1.

**Table 1:** Glossary of terms and abbreviations. Term Meaning.

| Term | Meaning |
|---|---|
| **Spike sorting and evaluation** |  |
| **KS4, IC, MS5, SC2** | Kilosort4, IronClust, MountainSort5, SpyKING CIRCUS2 |
| **TP, FN, FP** | True positive, false negative, false positive |
| **TPR, FNR, FDR** | True-positive rate (recall), false-negative rate, and false-discovery rate; precision = 1 − FDR |
| **F1** | The harmonic mean of precision and recall |
| **NM (no merge)** | The single detected unit best matching the ground-truth neuron |
| **BM (best merge)** | Units merged in descending order of precision, stopping when F1 first falls |
| **Spike and waveform properties** |  |
| **Isolated spike** | A spike whose preceding interspike interval exceeds 1 s |
| **Relative height** | A spike’s amplitude divided by the median amplitude of nearby isolated spikes |
| **Attenuated** | Relative height below the 5th percentile of the isolated spikes’ relative heights |
| **Full-height** | Not attenuated |
| **Burst number** | Position within a run of spikes separated by ISIs of at most 5 ms |
| **Residual magnitude** | Norm of the recorded minus the predicted extracellular waveform |
| **Optimal threshold** | Residual magnitude best separating clean spikes from simulated overlaps |
| **Above / below** | Residual magnitude above or below the optimal threshold |
| **Clean / coincident** | No spike of another unit within ±1 ms, or one or more |
| **Matched** | Distance to the nearest predicted waveform below the 95th percentile for clean ground-truth spikes |
| **Error categories and analyses** |  |
| **Similar-template FP** | An FP from a nearby neuron with a waveform resembling the ground-truth unit’s waveform |
| **Inferred-overlap FP** | An FP from the sorter wrongly attributing residual voltage during a putative overlap |
| **Excess FN rate** | FNs above the threshold beyond the below-threshold rate, divided by the number of ground-truth spikes |
| **Excess FN fraction** | FNs above the threshold beyond the below-threshold rate, divided by the number of FNs |
| **Re-sort** | The sort obtained after subtracting the ground-truth neuron’s predicted waveforms |
| **Mislabeled FN** | An FN the sorter detected but assigned to another unit |

### Experimental methods

#### Animals and habituation

All animal work was approved by Virginia Tech IACUC. Ground-truth data were obtained from neocortex, thalamus, CA1, and subiculum using silicon–juxtacellular hybrid probes in awake head-fixed C57BL/6J mice (PND 42–150) navigating a 1-D treadmill (English et al., 2017). Briefly, mice were chronically implanted with titanium headplates affixed to the skull overlying cerebellum, and also with a stainless-steel ground wire parallel to the surface of the cerebellum.

#### Selection of brain areas

Neocortex, hippocampus, and thalamus were chosen as recording areas for three reasons: (1) these areas are commonly recorded by many users of spike sorting; (2) each area has a different population distribution of firing rates, cell sizes, and soma densities, all of which are hypothesized to affect spike sorting; and (3) these areas can be accessed from a single craniotomy (−1.75 posterior / 1.75 lateral, in mm from bregma). Recording from multiple areas in the same animal reduces animal numbers and increases efficiency. Although different areas could be accessed in the same animal with multiple craniotomies, we found that the stability and duration of recordings were substantially decreased if more than one craniotomy was made per hemisphere.

#### Production of silicon–juxtacellular hybrid probes

Silicon–juxtacellular hybrid probes were produced as described in English et al. (2017). Briefly, juxtacellular electrodes were pulled from 1 mm OD / 0.7 mm ID borosilicate glass and had impedances of 8–10 MΩ when filled with 130 mM potassium acetate. These juxtacellular electrodes were glued to 32-channel silicon probes (A1-32-Poly3, NeuroNexus) using light-cure dental acrylic. Using the inter-site spacing of the silicon probe as a visual distance guide, the tip of the juxtacellular electrode was fixed at a distance of ∼20 *µ*m from the nearest silicon probe recording site. The closest site was selected as a site in the middle (top to bottom) of an outer column of recording sites. The close distance between the juxtacellular tip and the recording sites makes it likely that both devices record the same neuron and that the neuron’s waveform on the silicon probe has sufficient signal-to-noise ratio (SNR) for spike sorting (see Table S1 for juxtacellular and extracellular SNRs).

#### Recording ground-truth data in awake behaving mice

Mice were placed in the head-fixed treadmill apparatus, the protective silicone elastomer was removed from the cran-iotomy, the juxtacellular–silicon hybrid probe was inserted 0.1 mm into the brain, and the craniotomy was covered with silicone fluid to prevent drying and improve stability. The probe was advanced into the brain in ∼1–2 *µ*m steps until action potentials were observed on the juxtacellular electrode, at which point movement was ceased and data collection began. In a typical recording session, ∼4–10 juxtacellular neurons were obtained sequentially in a single axial penetration, usually yielding ∼1–2 neurons from each of the targeted structures.

#### Data acquisition

Electrical signals from the juxtacellular electrode were recorded and amplified through an analog microelectrode amplifier (Cygnus IR-183), then digitized via an Intan RHD2000 system. Signals from the silicon probe were amplified and digitized through the same RHD2000 system, which results in hardware synchronization of the two signals. Both signals were digitized at 30 kHz. By converting the digital signals in the RHD2000 to analog voltage and outputting them to a digital microprocessor (CED Power1401), we calculated online the juxtacellular spike-triggered average of each channel of the extracellular probe, and discarded recordings in which the triggered average did not contain a spike waveform on any channel. This drastically increases the rate of data collection, since once such a recording is discarded the electrode can simply be advanced further into the brain to obtain another recording.

#### Dataset and assignment of recording location

Each neuron is referred to by its recording region and an index (CA1-1 to CA1-6, NEO-1 and NEO-2, SUB-1 to SUB-3, and THA-1 to THA-5; Table S1), and the publicly released sessions are named accordingly. Recording locations were determined by a combination of stereotaxic coordinates and electroanatomy (e.g., the presence of sharp-wave ripples indicates electrode placement in the CA1 pyramidal layer). Post hoc visualization of recording tracks was not attempted due to multiple recording attempts per animal.

### Soft labeling of juxtacellular spikes

To quantify uncertainty in the juxtacellular recording, we construct a generative classifier for spike versus noise voltage snippets: class-conditional densities for candidate spike and noise events are fit separately and then plugged into Bayes’ rule (Murphy, 2023) to obtain posterior probabilities that each candidate juxtacellular event is a spike or noise (Fig. S1). This amounts to spike sorting on a single channel, but with an explicit model for noise. Since the juxtacellular recording will be used as “soft ground truth” for performance evaluation of extracellular spike sorting downstream, a robust noise model is emphasized: we only want to judge extracellular spike sorting performance to the degree of confidence that the juxtacellular recording provides.

Briefly, the juxtacellular signal was high-pass filtered and normalized by a robust estimate of its noise level. Spike templates were learned by clustering snippets sampled evenly across the amplitude range (so that small spikes are not drowned out by noise) and were used for template matching to align candidate events. The spike density was modeled as a Bayesian Gaussian mixture fit in a 4-dimensional PCA space of sampled juxtacellular voltage snippets, retaining only components whose *centroids* were spike-like. The noise density was modeled as an isotropic multivariate *t* distribution, whose heavy tails place more likelihood on noise near the spike–noise boundary and so make the labeling conservative. For a snippet ***J****_t_*, the posterior probability of a spike at time *t* is *P* (*signal* | ***J****_t_*) ∝ *P* (***J****_t_* | *signal*) *P* (*signal*) normalized over the two classes (signal and noise), with the prior *P* (*signal*) set, in a preliminary step, to the fraction of the recording occupied by spike snippets. We chose a generative rather than a discriminative classifier because labeled data are scarce and only approximate: generative classifiers approach their asymptotic error with fewer training examples (Ng and Jordan, 2001), and they yield a posterior over classes rather than only a decision boundary. To fit and calibrate the posterior model without any labeled data, we take a geometric approach (Fig. S1; Supplementary Methods).

### Performance evaluation

#### Surrogate ground-truth spike trains

To propagate juxtacellular uncertainty into downstream analyses, we sample from the posterior soft labels *p_t_* = *P* (*signal* | ***J****_t_*), *t* = 0*, …, N* − 1 (the *posterior trace*), as follows. The posterior is evaluated at candidate event times (template-match peaks), and *p_t_* = 0 elsewhere. Define the random spike train

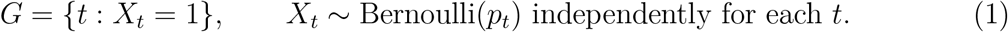

For each session we draw *K* = 1000 surrogate ground-truth spike trains *G*^(1)^*, …, G*^(^*^K^*^)^ as independent samples of *G*, indexed by *k*. Every performance metric below is computed separately on each *G*^(^*^k^*^)^, giving *K* values per session per metric. The random seed was fixed so that the same surrogates are used for every sorter, making the comparison across sorters paired at the level of the sampled labeling.

We report two summaries of the variability. The *95% interval across surrogates* conveys how uncertainty in the juxtacellular labeling affects the result: for each surrogate, we compute the median performance across neurons, and report the 2.5th and 97.5th percentiles of these *K* medians. The *interquartile range across sessions* conveys how much performance varies from neuron to neuron: for each neuron, we compute its median performance across surrogates, and report the 25th and 75th percentiles of these neurons’ medians. Reported point estimates are the median of the *K* per-surrogate medians.

#### Performance metrics

Let *D* be the set of detected extracellular spike times assigned to a ground-truth neuron, and let *δ* ≈ 0.83 ms be an empirically derived time delay tolerance (Supplementary Text). For each surrogate *G*^(^*^k^*^)^, a ground-truth spike is a *true positive* (TP) if at least one detected spike lies within *δ* of it, and a *false negative* (FN) otherwise. A detected spike is a *false positive* (FP) if no ground-truth spike lies within *δ* of it. TP, FN, and FP are thus labels on individual spikes. Writing 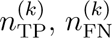, and 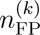 for the number of spikes carrying each label in surrogate *k*,

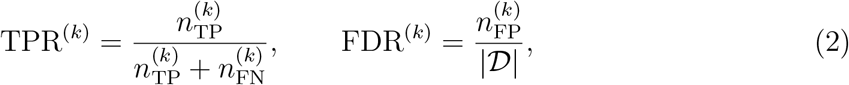

with FNR = 1− TPR, precision = 1− FDR, and F1 the harmonic mean of precision and recall (TPR). All metrics are computed separately for each surrogate, so each carries a distribution over the *K* sampled labelings rather than a single value (formal definitions in Supplementary Methods).

##### Extracellular SNR

Following Magland et al. (2020), each extracellular channel was bandpass filtered (300–6000 Hz, frequency-domain filter with error-function roll-offs), the mean waveform was computed at the ground-truth spike times, and SNR was its peak absolute amplitude divided by the noise level (median absolute deviation divided by 0.6745) on the channel where the peak occurred. SNR was computed for each of the *K* = 1000 surrogate ground-truth spike trains and summarized by the median.

#### Correlograms

For each surrogate, the auto-correlogram (ACG) of the ground-truth spike train and its cross-correlograms (CCGs) with the TP, FN, and FP events were computed as conditional rates: the rate of events at each lag relative to a ground-truth spike. Spike times were binned at 0.1 ms, lags up to ±25 ms were used, and the zero-lag bin was excluded. Average CCGs with uncertainty bands were computed across surrogates (Supplementary Methods).

#### Merge conditions

In spike sorting evaluation, one must determine which detected extracellular unit corresponds with the ground-truth unit; this problem might be non-trivial if there is a high degree of interneuronal synchrony, particularly if one wishes to account for errors arising from failures of merging and splitting clusters. We take two approaches.

- **NM (no merge).** Following Pachitariu et al. (2024), we match the juxtacellular neuron with the single detected unit that maximizes 1 − FNR − FDR, without any additional curation.
- **BM (best merge).** We rank all detected units by precision (1 − FDR) in descending order and then cumulatively merge units, stopping when adding the next unit first lowers F1. BM approximates the best merge attainable given the units the sorter detected, provided that a neuron’s oversplit clusters have higher precision than any other unit; whether they do is a hypothesis we test.

Both criteria are evaluated on the mean of each metric across the *K* surrogates, so each session’s NM unit and BM unit are chosen once and then held fixed in subsequent analyses, as repeating the selection for every surrogate would have been computationally prohibitive. We do not consider any additional strategies for splitting.

### Spike sorting

All four sorters were run on the same 16 sessions and evaluated against the same soft ground truth, with the same tolerance *δ*, the same *K* = 1000 surrogates, and the same NM and BM merge conditions, so per-session metrics are directly comparable across sorters.

#### KS4

KS4 (v4.0.24; Pachitariu et al., 2024) was run with its default settings. Drift correction was disabled (`nblocks` = 0), as recommended for a probe of this size. For the drift comparison, KS4 was run a second time with rigid drift correction enabled (`nblocks` = 1) and all other settings unchanged. To ask whether default parameters underestimated KS4’s performance, its parameters were also tuned by Bayesian optimization (hyperopt, tree-structured Parzen estimator; `acg_threshold, ccg_threshold, Th_universal, Th_learned, dminx, min_template_size, nearest_templates, and nearest_chans`; drift correction off). Optimization maximized the F1 score of the best-matching unit against the ground truth, with four-fold cross-validation across sessions (folds of four sessions): each session was sorted with parameters optimized on the other folds, using 30 evaluations per fold, each scored on four training sessions. Results in the main text use default parameters, as most users would.

#### IC

IronClust (IC; v5.9.8) was accessed through SpikeInterface (v0.103.2). Following the KS4 benchmark (Pachitariu et al., 2024), which reported that IronClust’s default parameters performed best, IC was run with all parameters at their defaults: detection threshold 3.5, adjacency radius 50 *µ*m, 300–8000 Hz bandpass filter, and trimmed-mean common referencing. IC clusters spikes by density-peak clustering (Rodriguez and Laio, 2014) on a *k*-nearest-neighbor graph, then merges clusters with similar waveforms; it does not use template de-convolution. Drift handling is part of clustering rather than a separate correction step: each spike’s neighbors are drawn only from time batches (300 s by default) with similar activity across the probe. This was left at its default, so unlike the other three sorters, drift handling was not disabled for IC. IC has no dedicated publication; see Jun et al. (2017) for JRCLUST, its predecessor.

#### MS5

MS5 was accessed through SpikeInterface (Buccino et al., 2020) and required parameter adjustment from its defaults before it clustered these recordings sensibly. The detection threshold was reduced from 6 to 4, the adjacency radius was reduced from 150 *µ*m to 45 *µ*m, and pre-whitening was enabled. With default settings MS5 severely under-clustered on the compact 32-channel Poly3 geometry used here, failing to isolate the ground-truth unit in several sessions. MS5 has no dedicated publication; see Chung et al. (2017) for the original MountainSort algorithm, including ISO-SPLIT clustering.

#### SC2

SC2 was accessed through SpikeInterface and run with the following deviations from the SpikeInterface defaults: detection threshold 6, high-pass cutoff 300 Hz, and motion correction disabled. SC2 has no dedicated publication; see Yger et al. (2018) for the original SpyKING CIRCUS. SC2 performs its own internal preprocessing; clustering is density-based and spike assignment uses Circus-OMP template matching.

### Errors associated with neuron-intrinsic variability

#### Juxtacellular waveform features

For the analyses below, the juxtacellular trace was high-pass filtered (100 Hz, third-order zero-phase Butterworth), and a spike’s peak time was taken as the maximum of that trace within ±0.5 ms of the spike time. Spikes were taken from the surrogate spike trains. To control for instability or drift in the juxtacellular channel and isolate physiological changes, for each spike, amplitude was expressed as a percentage of the largest-amplitude waveform occurring within one second of it (percent local max amplitude), a window short enough that the local maximum is a nearby spike, and events were binned into 21 equally spaced bins between 0% and 100%.

##### Spike attenuation

As another measure of attenuation relative to the neuron’s full-height spikes, amplitude was expressed relative to its *isolated* spikes, those preceded by more than 1 s of silence. Each spike’s *relative height*was defined as its amplitude divided by the median amplitude of isolated spikes within ±5 s. A spike was *attenuated* if its relative height fell below the 5th percentile of the isolated spikes’ relative heights. Relative height was then summarized as a function of the preceding ISI, in 24 logarithmically spaced bins from 1 ms to 2 s. The recovery time was the shortest preceding ISI at which the bin’s median relative height reached 95% of the isolated spikes’ median. Excess FNs were defined as (*n*_FN,att_ − FNR_full_ *n*_att_)*/n*_FN_, where *n*_att_ is the number of attenuated spikes and *n*_FN,att_ is the number of FNs among them, and FNR_full_ is the FNR of full-height spikes; negative estimates are plotted at zero but reported in the text.

#### CCG burst measures

Two features were extracted from the TP and FN CCGs: the lag of the peak, and the *CCG asymmetry*, defined as the CCG mass at positive lags minus the mass at negative lags, divided by the total mass. Asymmetry lies between −1 and 1; a positive value indicates more of the CCG’s mass lies above zero lag.

#### Interspike interval and burst number

Error was computed as a function of the interspike interval preceding each ground-truth spike, and separately as a function of burst number, where burst number *j* denotes the *j*-th spike in a run of consecutive spikes separated by interspike intervals of at most 5 ms. Positions beyond 8 were pooled into position 8.

Because the ground-truth spike train is probabilistic, burst position cannot be assigned to a fixed set of spikes. Conditional recall at each burst position was therefore estimated across surrogates: burst positions were labeled within each surrogate, and TP and total spike counts at each position were summed across all *K* surrogates before dividing. Per-session Spearman correlations between FNR and burst number were tested by exact permutation test. One session contributed fewer than three burst positions and was excluded, leaving 15 sessions. The pooled posterior mean FNR curve averages the per-session estimates with equal weight per session, so that neurons contributing many burst spikes do not dominate the pooled curve.

### Errors associated with extrinsic variability

#### LFP spectrum

The LFP power spectrum was computed in a 100 ms window centered on each ground-truth or detected spike. Logistic regressions were fit per session to predict TP versus FN and, separately, TP versus FP, with a 90/10 train-test split. Regressions were L1-regularized, with inverse regularization strength *C* = 0.1. Let *β_i_* denote the fitted logistic-regression coefficient for the spectral feature at frequency *f_i_*, for *i* = 1*, …, n_f_* with *n_f_* = 60 frequencies spaced logarithmically from 5 to 1000 Hz. For each fitted regression, the absolute-coefficient-weighted frequency center and spectral spread were calculated as

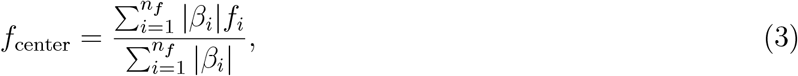

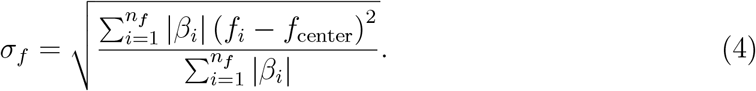

These quantities were calculated separately for each session and classification task and summarized across sessions. Above-chance prediction and the difference in AUC between TP-versus-FP and TP-versus-FN were assessed with Holm-adjusted tests, the latter by a one-sided exact paired session-level sign-flip test.

#### Probe and brain motion

KS4’s drift correction is not recommended for a probe of this geometry. As a sensitivity check, KS4 was additionally run with drift correction enabled and the two runs compared (Fig. S5D). Errors were also examined as a function of raw spike depth (Fig. S5C), and the mean absolute spatial distance per session was computed between true positives and false negatives, and between true positives and false positives, under both conditions (Fig. S5E).

#### Spike overlap and burst-related waveform variation

##### Predicted waveforms

All analyses in this section used the NM unit of the drift-off KS4 sort. For each neuron, the extracellular waveform of every ground-truth spike was predicted from its juxtacellular waveform with a multi-task LASSO, cross-fitted with 5-fold cross-validation so that each spike’s predicted waveform came from a model fit without that spike. Because refitting the LASSO for every surrogate spike train was not computationally tractable, ground-truth spikes used to fit the LASSO were defined once per session by thresholding their posteriors at 0.5, and the resulting out-of-fold predictions were stored. Because very few candidate events had intermediate posterior probabilities (Fig. S2), the results should not be highly sensitive to this threshold and all analyses that use the predicted extracellular waveforms followed this convention. Extracellular waveforms were taken from a recording that was high-pass filtered (300 Hz, fourth-order zero-phase Butterworth) and ZCA-whitened in 10-second chunks, each with its own covariance (regularized by 10^−5^). Within each training fold, let **X**_train_ ∈ ℝ*^n^*^train×61^ hold the standardized juxtacellular wave-forms of the training spikes (61 samples), with **X**_train_*_,j_* ∈ ℝ*^n^*^train^ its *j*-th column (one jux-tacellular time sample), and **Y**_train_ ∈ ℝ^*n*_train_×1952^ their standardized extracellular waveforms (32 channels × 61 samples). The penalty was set relative to the data, so that regularization had comparable strength across sessions and folds: *α* = 0.01 *α*_max_, where

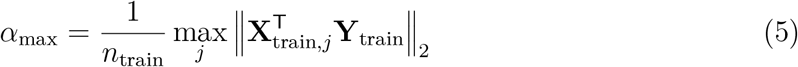

is the smallest penalty for which all coefficients are zero.

##### Waveform alignment

Small differences in spike timing can make two otherwise similar waveforms appear different, so all comparisons require a consistent alignment. Extracellular waveforms of TPs and FNs were extracted in ∼2-ms windows (20 samples before to 40 after) aligned to their ground-truth spike times. FPs have no juxtacellular spike, so their extracellular waveforms were extracted at KS4 detection times, shifted by a per-session offset to match the ground-truth convention (the median lag between each TP’s juxtacellular spike time and its matched KS4 detection). TPs and FNs were compared with FPs directly. Because the offset aligns the two conventions only on average, FP distances may be slightly inflated relative to TP and FN distances, which would affect all FP comparisons alike.

##### Distance to predicted waveforms and matching

For each event type, we computed the Euclidean distance between its recorded extracellular waveform (32 channels × 61 sam-ples, whitened, aligned as above) and the nearest out-of-fold LASSO prediction of any ground-truth spike in the session. For TPs and FNs, the spike’s own prediction was excluded, so that every event is compared only with predictions of other spikes, as an FP necessarily is; FPs have no prediction of their own, but we wish to compare them to the distribution of predictions. As a baseline noise level for the neuron’s waveform variability in the absence of overlaps, we considered the distribution of such distances for *clean* ground-truth spikes (those with no spike of another KS4 unit within ±1 ms, excluding putative oversplit clusters of the ground-truth neuron; precision > 0.5). An event of any type was defined as *matched* if its distance fell below the 95th percentile of these clean ground-truth distances. To prevent a clean spike from setting its own baseline, the clean ground-truth spikes were split at random into two and each half was judged against the threshold from the other; all other events (coincident ground-truth spikes and all FPs) were judged against the thresh-old from all clean ground-truth spikes. Because the threshold is the 95th percentile of the clean ground-truth distances, the matched fraction of clean ground-truth spikes is 0.95 in expectation, owing to the cross-fitting. The matched fraction of a group is the proportion of its events that were matched, computed for groups of at least 10 events. Because the matched fraction falls as coincidences rise, coincident TPs and FNs were weighted to the FPs’ distribution of nearby-spike counts (1, 2, or 3 or more) before comparison with coincident FPs. This way, differences in matched fraction between FPs and the other event types reflect how much their waveforms resemble the neuron’s predicted waveforms, regardless of the coincidence rate.

##### KS4 spikes near each error type

We counted the KS4 spikes within ±1 ms of each TP, FN, and FP in KS4’s time convention, counting all units other than potential oversplit units of the ground-truth neuron as before. The NM unit’s own detection, where present, is included once (more on this later). For each neuron, these counts were averaged within each error type. The same counts were computed in the re-sort (below), using the re-sort units and the same exclusion of oversplit units. Groups were compared within each sort by two-sided sign tests.

##### Clean and coincident events

Let an event be defined as *coincident* if a spike of any KS4 unit other than the NM unit (again excluding putative oversplit units of the ground-truth neuron) falls within ±1 ms of it, and *clean* otherwise. This terminology will be used in various places.

##### Fraction of FPs attributable to inferred overlaps

For FPs, let FDR_clean_ and FDR_coinc_ be the false discovery rates among clean and coincident detections of the NM unit, *N*_coinc_ the number of coincident detections, and *N*_FP_ the number of FPs. The estimated fraction of FPs attributable to inferred overlaps is

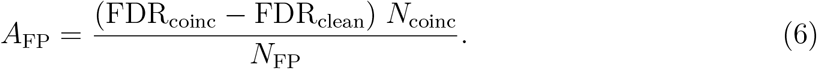

FDR_clean_ is the rate at which the NM unit produces FPs when no other unit fires nearby, and *A*_FP_ estimates the FPs among coincident detections in excess of that rate, as a fraction of all FPs (the reasoning is detailed in the Results). This assumes that the FPs unrelated to overlaps occur at the same rate among coincident as among clean detections; a negative estimate indicates that the assumption did not hold for that neuron, or reflects sampling variability, and is plotted at zero yet reported explicitly in the text in the few cases this happened.

##### Merging the next-cleanest unit

To ask whether FNs could be recovered by merging, we took, for each neuron, the KS4 unit with the highest precision against the ground truth other than the NM unit, and recomputed recall, precision, and F1 after merging it into the NM unit. This is similar to the precision ranking used by the best-merge (BM) procedure described above, but applied as a single merge into the NM unit, perhaps more analogous to how users of KS4 might perform a merge under the assumption oversplit clusters would have the highest precisions. As a check on this choice, we also applied KS4’s own refractory criterion between each candidate and the NM unit: a candidate was refractory if the number of its detections within ±1 ms of an NM detection fell below 0.25 times the number expected from the 20–50 ms shoulders of their CCG.

##### Subtraction and re-sorting

To ask whether FNs had been missed outright or assigned to other units (i.e., mislabeled), we artificially removed the ground-truth neuron from each recording and sorted it again. We call this sort the *re-sort*. For each ground-truth spike, its LASSO-predicted extracellular waveform was mapped from whitened space back to data space and subtracted from the raw recording at the ground-truth spike’s time. For this mapping, the inverse of each chunk’s whitening matrix was computed exactly and applied to the predictions falling in that chunk. KS4 (v4.0.24) was then rerun on the resulting recordings with the same settings as the default drift-off sort (`nblocks` = 0).

For each ground-truth spike, we counted the KS4 detections of any unit, including pu-tative oversplit clusters, within ±1 ms before and after subtraction. We say an event *lost a detection* if its count decreased by at least one. A mislabeled FN is a detection of the ground-truth spike under another unit, so it should disappear when the neuron is removed, as a TP’s detection does, whereas a spike missed outright leaves nothing to remove (i.e., nothing that the sorter noticed in the original sort). Within attenuated and full-height spikes as separate groups (as defined for earlier), the mislabeled fraction was estimated as

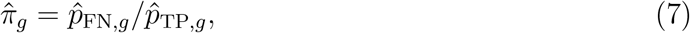

where *p̂*_FN_*_,g_* and *p̂*_TP_*_,g_* are the fractions of FNs and TPs in group *g* that lost a detection; dividing by the TPs’ fraction corrects for imperfect subtraction. Groups with at least 10 TPs and 10 FNs and *p̂*_TP_*_,g_ >* 0.2 were included and combined weighted by their FN counts, 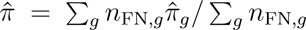. As a control for changes the re-sort makes elsewhere in the recording, the fraction losing a detection was also computed at random times sampled at least 5 ms from any ground-truth spike (five times as many samples as there were ground-truth spikes; Fig. 5H, Supplementary Text).

##### Chance recall

To verify that the ground-truth waveforms were successfully subtracted (i.e., from the perspective of the sorter), we compared the recall of the re-sort’s best-matching unit with the recall expected from a uniform random clustering of all detections in that sort.

If the subtraction was successful, this recall should be consistent with that of a random clustering of the re-sort’s detections. The re-sort’s unit labels were shuffled across its spikes 1000 times, preserving every unit’s size, and the best-matching unit (the maximum of 1 − FNR − FDR across units) on each shuffle was scored against the ground truth. The same was done for the original sort, for reference. This analysis also provides evidence that all our units had sufficiently high SNRs for the sorter to perform inference on them. The chance level is the median shuffled recall across neurons, with a 95% interval across shuffles.

##### Residuals and simulated spike overlaps

For each ground-truth spike, the *residual* was its recorded waveform minus its predicted waveform, and the *residual magnitude* the Euclidean norm of this difference. To obtain a rough estimate of the residual magnitudes characteristic of spike overlaps, *clean ground-truth spikes* (TPs and FNs with no spike of another KS4 unit within ±1 ms, oversplit units excluded) were used to simulate overlaps: 10,000 per neuron, each the sum of a clean ground-truth spike’s extracellular waveform and the mean waveform of another unit at a lag drawn uniformly from ±1 ms, with residuals computed against the clean spike’s predicted waveform. The *optimal threshold* was the residual magnitude maximizing Youden’s *J* = sensitivity + specificity −1, where sensitivity is the fraction of simulated overlaps above the threshold and specificity the fraction of clean ground-truth spikes below it.

##### Excess FN rate

Ground-truth spikes were classified as *below* or *above* by whether their residual magnitude fell below or above the optimal threshold. Analogously to *A*_FP_ above, let *n*_FN,above_ − FNR_below_ *n*_above_ be the number of FNs above the threshold in excess of what the below-threshold FNR would predict: an estimate of the number of FNs that spike overlaps could plausibly account for. The *excess FN rate* is this number per ground-truth spike,

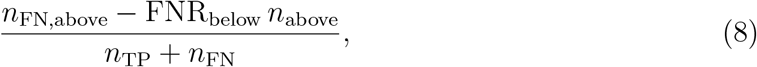

and the *excess FN fraction* is the same number divided by *n*_FN_ instead, i.e., the rate divided by FNR; neither is floored. Because the optimal threshold separates clean spikes from simulated overlaps imperfectly, both are likely underestimates, and we interpret them primarily by how other quantities co-vary with them. Principal components, computed from predicted and ground-truth waveforms, were used for visualization only.

##### Intrinsic and extrinsic waveform variation

For the predictive comparison, sessions were retained if the cross-fitted model improved median held-out waveform mean-squared error relative to the training-fold mean waveform (13 of 16 sessions); analyses based on distance and residual magnitude used all neurons. Ground-truth spikes preceded by more than 200 ms of silence formed the *reference set*, and the reference waveform **r** was the mean predicted waveform of reference-set training events within each classifier training fold. For event *i* with extracellular waveform **x***_i_* and predicted waveform **x̂***_i_*,

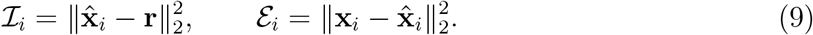

**I***_i_*, the deviation of a predicted waveform from the reference waveform, is variation the juxtacellular channel predicts, which we call *intrinsic*; *ε_i_*, the deviation of the recorded waveform from its prediction, is variation it does not, which we call *extrinsic*. Burst-related waveform change contributes to the intrinsic term (Fig. 6E), spike overlaps, recording noise and model error contribute to the extrinsic term, and motion of the neuron relative to the probe likely contributes to both, which we were unable to model further. Per-neuron Spearman correlations of preceding ISI with **I** and *ε* were tested by sign test across neurons. The contribution of each to held-out prediction of TP/FN status was compared by Shapley values (Supplementary Methods). Note that a less strict criterion than the 1 s used to define isolated spikes above was used here, trading conservatism for sample efficiency. Such bias would reduce the intrinsic term rather than inflate it.

##### FNR by attenuation and residual magnitude

Each neuron’s ground-truth spikes were classified as attenuated or full-height (by the juxtacellular waveform, as above) and as below or above the optimal threshold (see residuals and simulated spike overlaps, above), and the conditional FNR was computed in each of the resulting four groups. Each group’s fraction of a neuron’s FNs was then its number of spikes times its FNR, divided by the neuron’s total number of FNs. Differences between groups, and each group’s fraction of FNs relative to its fraction of spikes, were tested across neurons by two-sided sign tests.

To characterize sources of slow time-varying error (e.g., drift associated), each session’s ground-truth spikes were divided into ten equal intervals and recall computed in each interval with at least 20 spikes; the Spearman correlation between interval index and recall was tested by exact permutation. As a control for the four-group split described in the previous paragraph, the 30% of intervals with the lowest recall were removed within each neuron and the split recomputed on the remaining spikes. Removing intervals rather than whole sessions avoids confounding time-varying error with differences in overall FNR between neurons.

##### Sensitivity to rejected ground-truth spikes

To test whether FPs include ground-truth spikes the soft labeling rejected, all FP analyses were repeated after removing FPs within ±1 ms of any threshold crossing on a more permissive 5*σ* amplitude detector on the juxtacellular trace (Supplementary Text).

### Comparison across sorters

The burst analysis and the comparison of intrinsic and extrinsic waveform information were repeated for IC, MS5, and SC2 on the same 16 sessions (13 sessions with informative wave-form predictions for the information comparison). For each sorter the analysis was applied to the NM unit. For the burst analysis, the conditional recall at each burst position was computed exactly as for KS4, with the coverage mask built from that sorter’s detected spike times. For the information comparison, the cross-fitted waveform predictions depend only on the juxtacellular and extracellular data and are therefore identical across sorters; only the TP/FN classification differs.

Per-session Spearman correlations between FNR and burst position were computed for each sorter across the burst positions at which that session contributed data, with the same exclusion as for KS4. Statistical tests are described under Experimental Design and Statistical Analysis.

### Experimental Design and Statistical Analysis

The experimental unit is the ground-truth neuron; each of the 16 sessions contributed one juxtacellularly recorded neuron. Sessions were included if the online juxtacellular spike-triggered average contained a spike waveform on at least one extracellular channel. No statistical method was used to predetermine sample size, which was limited by the yield of awake juxtacellular recording. Ground-truth uncertainty was propagated by computing every metric on *K* = 1000 surrogate spike trains (Surrogate ground-truth spike trains); we report medians with the 95% interval across surrogates and the interquartile range across sessions. Medians and interquartile ranges were used because they are robust to atypical values, which could arise in individual sessions from labeling errors, periods of noise, or declining neuronal health. Given the small sample and skewed distributions, all tests were nonparametric: permutation tests for group comparisons (group labels reassigned 10^4^ times), exact permutation tests for per-session Spearman correlations (*ρ_s_*), sign tests for per-session *ρ_s_* against zero (preferred to signed-rank tests because the *ρ_s_* distributions are left-skewed), and exact paired Wilcoxon signed-rank tests for within-session comparisons between sorters; LFP classifier performance was compared against chance and between error types with exact paired sign-flip tests. Comparisons in Figure 5 and Figure 6 used exact paired sign-flip permutation tests (all 2*^n^* sign combinations of per-neuron differences), with effect sizes reported as the median per-neuron difference; relations with FNR across neurons were tested with Spearman correlations. Comparisons of nearby-spike counts (Fig. 5F) and of fractions losing a detection (Fig. 5H) used two-sided sign tests. For comparisons between clean and coincident events, whose mean differences were dominated by a few neurons, the two-sided exact sign test on the per-neuron direction is reported alongside. Attributable fractions were not floored at zero; negative estimates are shown at zero and listed in the text. Tests were two-sided except where the hypothesis was directional: the LFP classifier comparisons (above chance, and TP-versus-FP predictability exceeding TP-versus-FN), the test that intrinsic waveform variation increases at short interspike intervals (*ρ_s_ <* 0), and the comparison of Shapley values (*ϕ**_I_** > ϕ_ε_*), which were one-sided. Multiple comparisons were corrected with the Benjamini–Hochberg procedure within each family of tests (18 between-sorter comparisons per merge condition: six sorter pairs × three metrics, F1, recall, and precision, of which only the F1 comparisons are discussed in the Results), and with the Holm procedure for the LFP classifier tests and for the Shapley comparisons across the four sorters. Exact *p* values are reported throughout, corrected where a correction applies and not further labeled in the text; results described as significant are those with *p <* 0.05 after correction where applicable.

### Data and code availability

Curated ground-truth data will be made available online at dandiarchive.org. Code for soft labeling the juxtacellular data is at: https://github.com/zach-saccomano/juxta-softlabel.

## Results

### Simultaneous juxtacellular and extracellular recording with soft ground-truth labeling

We recorded 16 single units juxtacellularly with simultaneous 32-channel silicon probe recording in neocortex, thalamus, CA1, and subiculum of awake mice (Fig. 1A). Although the juxtacellular signal has a much higher SNR than the extracellular signal (see Table S1), juxtacellular spike detection is nontrivial (Fig. S2). The difficulty is greatest for bursting neurons: even in high SNR recordings, small-amplitude events late in a burst are continuous with the noise rather than cleanly separable from it. In the PCA basis of the juxtacellular snippets, embedded juxtacellular waveforms form an elongated, curved distribution whose smallest-amplitude part overlaps the noise distribution (Fig. 1B; Fig. S1A). Any hard thresh-old is therefore problematic: too conservative, and true low-amplitude spikes are missed; too permissive, and many noise events are wrongly labeled as spikes.

**Figure 1:**
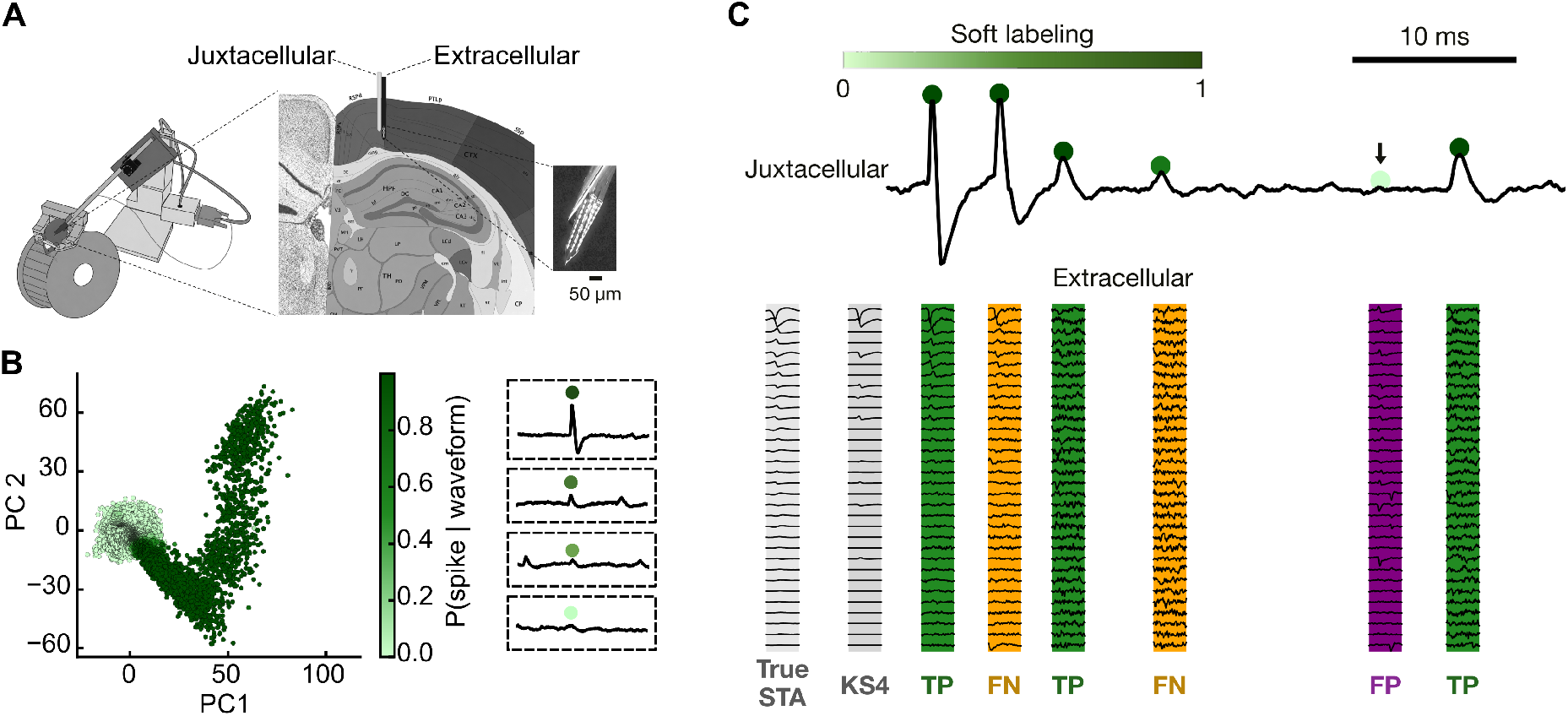
Robust performance evaluation of spike sorting in the awake mouse with juxtacellular ground truth. (A) Left, awake head-fixed mice on a running wheel. Right, single units were recorded juxtacellularly with simultaneous multi-channel extracellular recording in neo-cortex, CA1, subiculum, and thalamus. (B) Left, juxtacellular snippets from a neocortical unit in their PCA basis, colored by posterior probability of a spike. Right, example snippets spanning the range of posterior probabilities (Fig. S1). (C) Top, juxtacellular trace with candidate events and their soft labels (posterior probability of a spike; Fig. S1, Methods). The fourth event is uncertain (*p* = 0.66) and the fifth is almost certainly noise (*p <* 0.001; arrow) despite a sorter detection. Bottom, multi-channel extracellular waveforms for candidate events, beside the soft label-weighted spike-triggered average (STA) and KS4’s template for the best matched unit; channels are sorted by STA amplitude. Error labels (TP, FN, FP) are for one sampled ground-truth spike train.

We therefore modeled spike and noise snippets explicitly and combined the two densities through Bayes’ rule, assigning each prescreened candidate event a posterior probability of being a spike, which declines gradually toward the noise cloud rather than at a sharp boundary (Fig. S1F; Methods). Recall, precision, and F1 are then computed as posterior-weighted (‘soft’) quantities: each candidate event is included in a sampled ground truth spike train with probability equal to its posterior, like a weighted coin flip, and this is repeated 1000 times. In Fig. 1C, for example, the fourth event (posterior 0.66) appears in about two-thirds of the surrogate spike trains, events with posterior *>* 0.99 appear in almost all of them, and the fifth (posterior *<* 0.001) is judged almost certainly noise despite the sorter having detected a spike there. TPs, FNs, and FPs are determined for each surrogate, and we report medians and uncertainty intervals over the 1000 surrogates. Because our goal is to test the assumptions of spike sorting qualitatively rather than to provide well-calibrated error estimates, we designed the soft labeling to be deliberately conservative (Methods; Discussion).

### Overall performance of KS4

We characterize spike sorting error in depth for KS4, likely the most widely used sorter and a mature template deconvolution approach whose assumptions we wish to examine. A final section tests our main findings in other sorters. KS4 was run using its recommended settings for our probe (see Methods). We adopted the convention that a detected spike counts as ‘matching’ a ground-truth event if it falls within a tolerance *δ* of it. Performance metrics were insensitive to this exact choice: each session’s metrics saturated by 0.23 ms on average (s.d. 0.21 ms under NM, 0.23 ms under BM; at most 0.77 ms) and stayed saturated at larger tolerances, so we used *δ* ≈ 0.83 ms throughout which covered the worst case (Fig. S3B; Supplementary Text). Because the ground-truth unit may be split across several sorted units, we report two merge conditions: NM (no merge), where we took the single best matched extracellular unit the spike sorter produced without any additional curation, and BM (best merge) which uses ground-truth information to approximate the *best possible merge* that could be performed on cleanly oversplit clusters given the units the sorter detected (Methods). We illustrate the idea using CCGs between ground-truth spikes and either TP, FN, or FP extracellular spikes in an example session (Fig. 2A–C). In this example, a substantial number of ground-truth spikes fell in the false negative CCG under NM (Fig. 2A); whereas F1 peaked when units with the 7 highest precisions were merged (F1 = 0.687) (Fig. 2B), and mass shifted from the false negative to the true positive CCG as units were merged (Fig. 2C).

**Figure 2:**
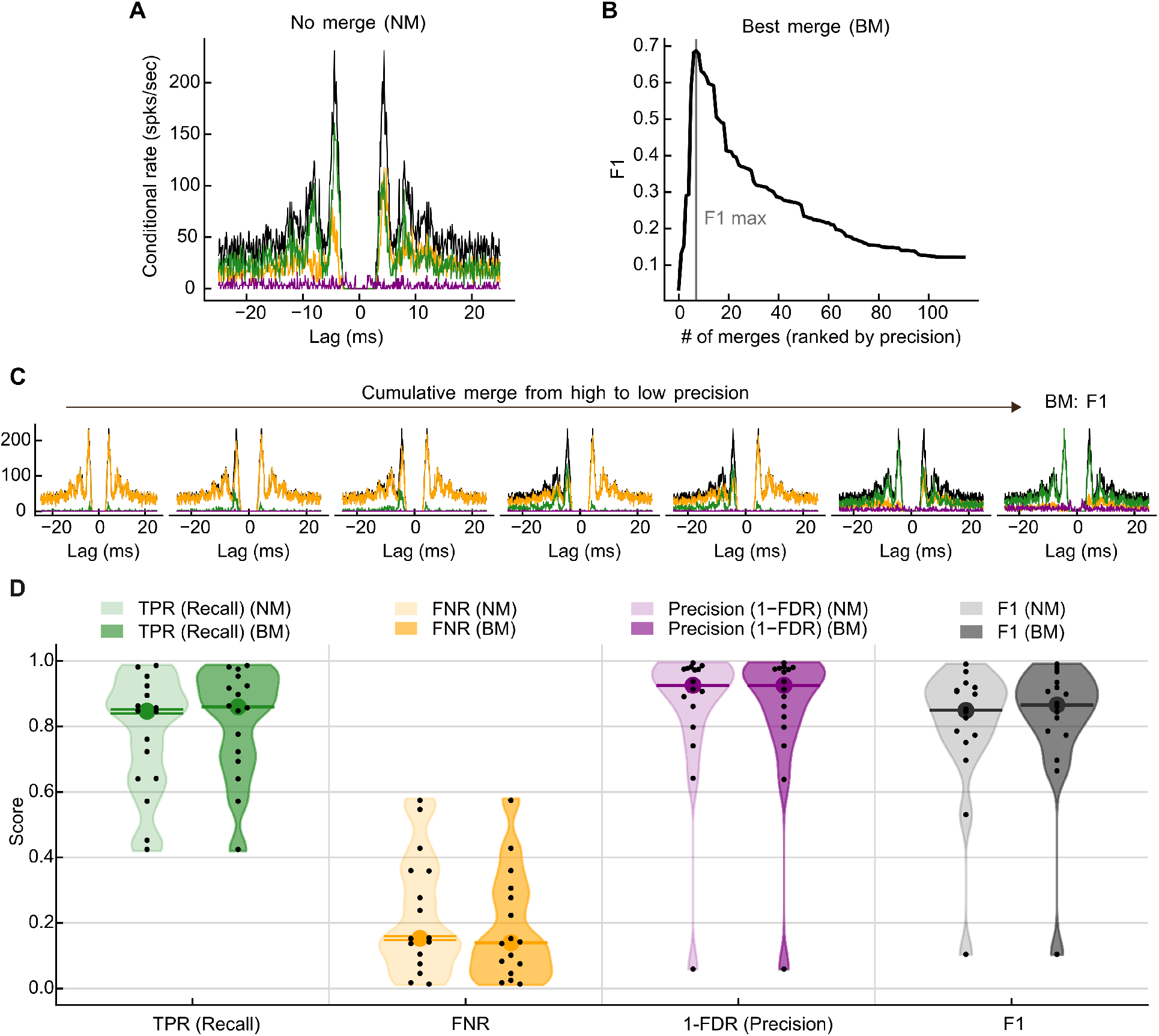
Overall recall is lower and more variable across recordings than precision. (A–C) Example session (NEO-2). (A) Posterior mean ACG of the ground-truth unit (black) and CCGs between the ground-truth unit and the TP (green), FN (orange), and FP (purple) spikes of the NM unit (Methods). (B) F1 of the cumulatively merged unit versus number of detected units merged, ranked from high to low precision; BM is the F1 maximum (gray line). (C) Posterior mean ACG and CCGs at each step of the cumulative merge, ending at BM. (D) Overall performance under NM (light) and BM (dark). Black dots, sessions (*n* = 16); colored dot and bars, median and 95% interval of the across-session median across juxtacellular posterior surrogates.

Across recorded ground-truth neurons, median NM recall was 0.847 whereas median NM precision was 0.927 (Fig. 2D). Furthermore, across ground-truth neurons, NM recall varied about twice as much as precision (Q1–Q3 of 0.642–0.903 for recall versus 0.846–0.976 for precision under NM), meaning that recall is both poorer and more variable across neurons than precision is. Without merging, KS4 performed best for thalamic units (*N* = 5 neurons, median F1 0.933, Q1–Q3 0.854–0.968), and weakest for those recorded in the neocortex (*N* = 2 neurons, median F1 0.653, Q1–Q3 0.592–0.713) (Fig. S4). Notably, the best merge (BM) condition only produced a modest improvement in overall performance (median F1 0.849 to 0.867).

For BM, since candidate units are merged in descending order of precision, a ground-truth unit cleanly oversplit into several high-precision clusters would be largely reassembled by this procedure. But that is not what we see. This suggests missing spikes are either absent from the sorter’s output entirely or present only in clusters each too imprecise to add without a net cost in F1; we will test this directly in Fig. 6G,H. This further suggests a hypothesis we pursue in the following sections: that error concentrates on the events for which the template is least valid. In particular, that recall depended on brain region and was highly variable across neurons, naturally suggests the deficit in recall depends on intrinsic properties of individual neurons, like bursting.

To test the sensitivity of these performance metric estimates in KS4, we additionally report all marginal metrics using two different ways of defining ground-truth (Fig. S3). Importantly, estimates of recall using our ground-truth labels were considerably higher than other labeling methods and estimates of precision only deviated slightly, consistent with the strategy discussed in the methods. These estimates were also robust to KS4’s parameters and to signal quality. Parameters tuned by cross-validated Bayesian optimization raised recall (0.868 vs. 0.847 with defaults; higher in 13 of 16 sessions) but not F1 (0.859 vs. 0.849; higher in 10 of 16 sessions, median paired difference 0.002, Wilcoxon signed-rank *p* = 0.71; Fig. S5A; Methods), so we report default parameters, as most users would run them. Performance showed no detectable relationship with extracellular SNR (median 4.1, IQR 3.5–5.4; *R*^2^ ≤ 0.01 for recall, FNR, precision, and F1 against log SNR; Fig. S5B). Under the strong assumption that a linear fit of each metric against SNR holds beyond the support of the data, moving each neuron along that fit to SNR = 8, SpikeForest’s inclusion cutoff (Magland et al., 2020), would raise median F1 only modestly, to 0.886 (Fig. S5A). Enabling drift correction, likewise, slightly raised recall but lowered precision and F1 (Fig. S5D; Table S2; Supplementary Text).

### Errors associated with neuron-intrinsic variability

As a next step, we quantified recall with respect to intrinsic properties of the ground-truth neuron, as measured on the juxtacellular channel (Fig. 3A). Juxtacellular spike amplitude was normalized relative to the largest spike in a one-second window to control for potential instability or drift in the juxtacellular recording and to isolate changes intrinsic to the neuron (Methods). Recall increased approximately linearly with percent local max amplitude of the juxtacellular spike (mean *R*^2^ of the per-session linear fit 0.69 ± 0.22; Fig. 3B). Because of the normalization, this already suggests that bursting is a major driver of FNR. As we will show later, this variation in the juxtacellular waveform was also present in the extracellular waveform and predicted which spikes are FNs (Fig. 6E,F). FNR also increased with burst number (position within a run of spikes separated by ISIs of at most 5 ms) in the pooled posterior mean (*ρ_s_* = 0.95, *p <* 0.0001; Fig. 3C); by exact permutation test, 6 of 15 sessions had *p <* 0.05.

**Figure 3:**
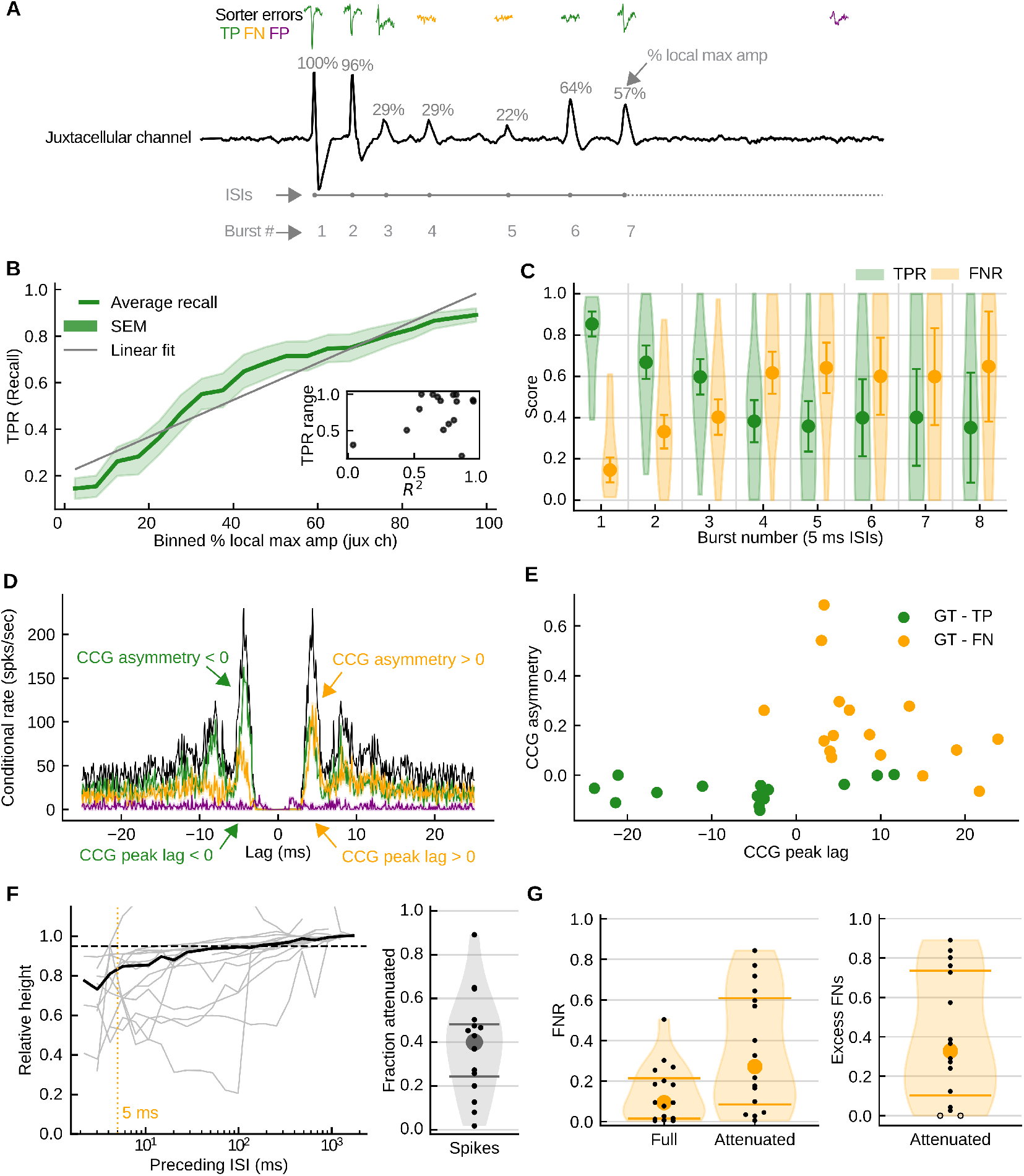
Sorting errors associated with neuron-intrinsic features. (A) Juxtacellular fea-tures (percent local max amplitude, preceding ISI, burst number) on an example trace with sorter errors (TP green, FN orange, FP purple). (B) Mean TPR versus binned percent local max amplitude; shading, SEM across sessions; gray line, linear fit; inset, per-session *R*^2^ and TPR range. (C) FNR and TPR versus burst number (ISI ≤ 5 ms). (D) Example CCGs between the ground-truth train and its TPs, FNs, and FPs. (E) CCG peak lag and asymmetry, GT vs. TP and GT vs. FN. (F)Left, spike height relative to isolated spikes (preceding ISI *>* 1 s) versus preceding ISI; gray, neurons; black, median; dashed, 95%; dotted, 5 ms. Right, fraction of spikes attenuated. (G) Left, FNR of full-height and attenuated spikes. Right, excess FNs among attenuated spikes as a fraction of FNs; hollow dots, negative estimates shown at zero. Dots, individual neurons. Definitions and tests in Methods; KS4 NM unit, drift-off sort; *n* = 16 neurons.

To avoid imposing a fixed ISI cutoff, posterior mean CCGs (Methods) were computed between the full ground truth (GT) spike train and either its TPs or its FNs (Fig. 3D; all neurons in Fig. S6). Across neurons, the CCG peak lag was 15 ms greater for GT-FN than for GT-TP (permutation test across sessions, *p <* 10^−5^), and CCG asymmetry (mass at positive minus negative lags, divided by the total) was 0.26 units greater (*p <* 10^−5^; Fig. 3E), indicating that FNs tend to follow other spikes at short latency.

Detectability likely tracks spike height rather than ISI per se, so we next examined burst-related waveform attenuation directly. In these data, we observed that spike height recovers quite slowly; relative to the typical neuron’s isolated spikes (preceded by more than 1 s of silence), juxtacellular spike height took a median of 85 ms to recover to 95% of its isolated level (IQR across neurons 32–350 ms), far beyond a conventional ∼5-ms burst index definition (Fig. 3F, left; Methods). Let a spike be termed *attenuated* if it is smaller than 95% of the neuron’s isolated spikes. In the median neuron, this 95% cutoff corresponded to a spike 69% of the typical isolated-spike height (IQR across neurons 61–81%). In the median neuron, 40% of spikes were attenuated (IQR across neurons 24–48%; Fig. 3F, right). Attenuated spikes had a higher FNR than full-height spikes in 14 of 16 neurons, and were at least doubled in 9 neurons (median FNR 0.27 vs. 0.09; IQR across neurons 0.09–0.61 vs. 0.02–0.21; Fig. 3G, left). FNs among attenuated spikes in excess of the full-height FNR amounted to a median of 4% of a neuron’s ground-truth spikes (IQR 1–10%), or a third of its FNs (IQR 10–74%; Fig. 3G, right). Two neurons had negative estimates, which we plotted at zero (CA1-2, −0.28, whose NM unit captured mainly attenuated spikes, leaving full-height spikes as FNs; THA-4, −0.11, with FNR *<* 1%). Across neurons, FNR fell with spike height in 14 of 16. We conclude that spike-history-dependent changes in spike height contribute substantially to the poor recall in Figure 2, robustly across burst definitions. Moreover, this error contribution cannot be easily explained away by low extracellular SNR (Fig. S5B).

### Errors associated with extrinsic variability

We term extrinsic sources of variability as factors in the recording environment rather than from the ground-truth neuron’s physiology: the LFP spectrum surrounding a spike, real or presumed motion of the probe relative to the brain, and real or presumed overlapping spikes from other neurons.

#### LFP spectrum

The LFP power spectrum was computed in a 100 ms window centered on each ground-truth spike or FP detection, and we searched for spectral features that could explain errors. Rather than performing hypothesis tests in high dimensions, we asked if error labels could be predicted by the surrounding LFP activity in held-out data. Separate L1-regularized logistic regressions were fit within each session to classify TP versus FN spikes and TP versus FP spikes using a 90/10 train-test split. Across 16 sessions, the mean test-set AUC was 0.642 ± 0.025 for TP versus FN and 0.745 ± 0.034 for TP versus FP (mean ± SEM). Both classifications performed above chance (TP versus FN: *p* = 1.83 × 10^−4^; TP versus FP: *p* = 4.58 × 10^−5^). Importantly, TP-versus-FP predictability exceeded TP-versus-FN predictability by a mean paired AUC difference of 0.104 (one-sided exact paired sign-flip test, *p* = 0.0021; Fig. 4). Thus, LFP spectral content was associated with both error types, but more strongly with false positives.

**Figure 4:**
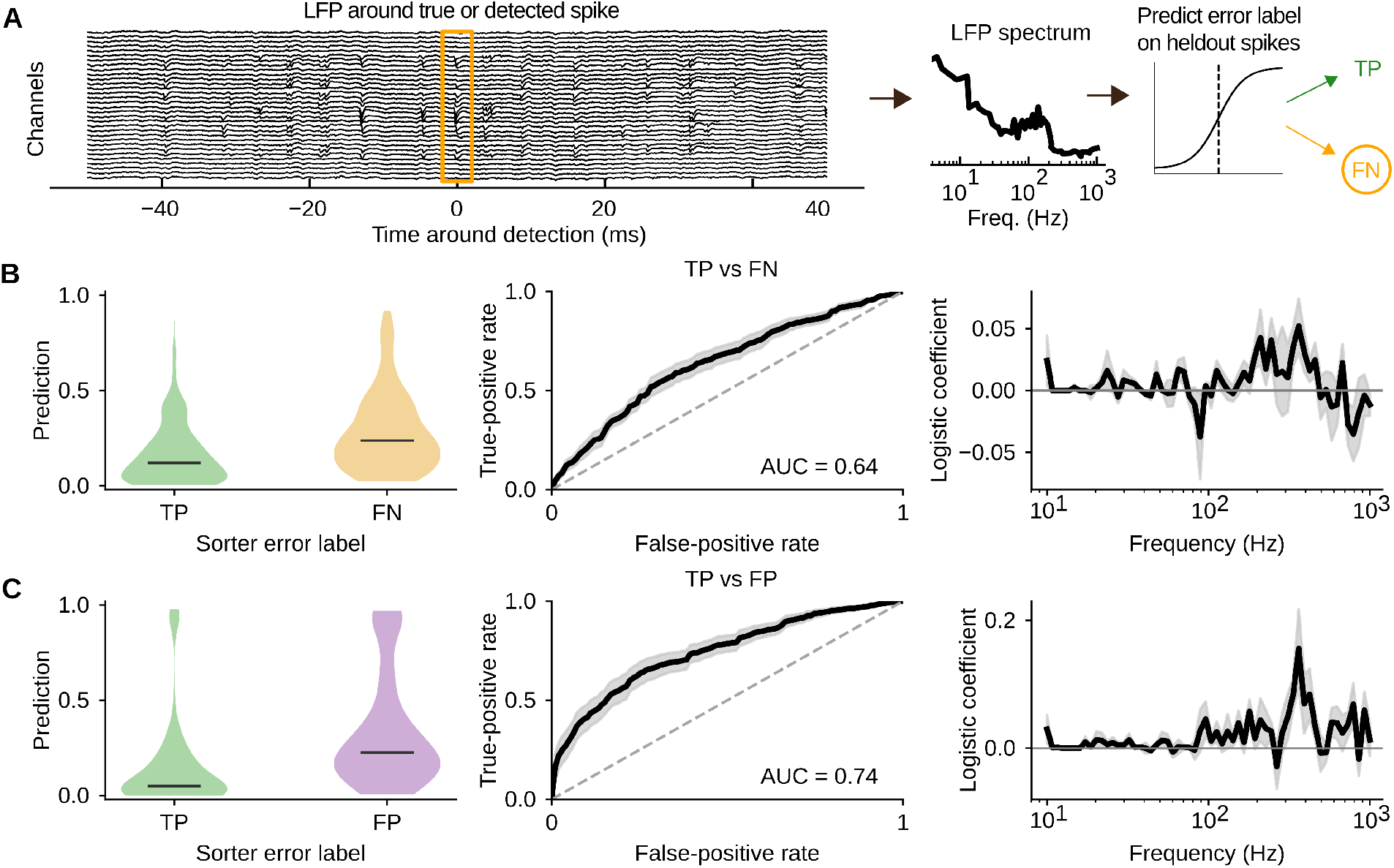
Sorting errors associated with the LFP spectrum. (A) LFP power spectrum in a 100-ms window centered on each ground-truth spike or FP detection (orange box); per-session L1-regularized logistic regressions classify TP versus FN and TP versus FP events on held-out data (Methods). (B) TP versus FN predictions. Left, held-out predictions grouped by error label (bars, medians). Middle, mean ROC curves across sessions. Right, mean logistic-regression coefficients across frequency. (C) Same for TP versus FP. Shading, SEM; KS4 NM unit, drift-off sort; *n* = 16 neurons.

Furthermore, the absolute-coefficient-weighted frequency center, a measure of where the coefficient magnitude is concentrated, was 377.9 ± 17.4 Hz for TP versus FP, with *σ_f_* = 250.3 ± 13.9 Hz, and 337.6 ± 21.0 Hz for TP versus FN, with *σ_f_* = 244.2 ± 18.4 Hz (mean ± SEM across sessions). Because this frequency range is consistent with elevated population spiking (377.9 Hz corresponds to a 2.64 ms period), these results raise the possibility that FPs are related to spike overlaps, which we examine momentarily.

#### Spike overlap

It has long been proposed that spike overlaps are a major challenge for spike sorting and that they might be resolved using algorithms like template deconvolution (Pillow et al., 2013). Such approaches assume that preprocessing and featurization yield stable templates, in the sense that oversplit clusters have sufficiently high precision that post hoc merging can reassemble them. With ground-truth data, we wish to avoid this assumption: we instead want to evaluate it. To this end, rather than assuming a stable template, we predict, for each ground-truth spike time, the extracellular waveform from the juxtacellular waveform with a cross-validated LASSO regression (Fig. 5A; Methods). This juxtacellular-predicted extracellular waveform can model variation from intrinsic factors (e.g., bursting), while absorbing extrinsic factors (e.g., putative spike overlaps) into the model’s residuals. In 13 of 16 sessions, cross-validated predictions reduced median held-out waveform mean-squared error relative to predicting the training-fold mean waveform, indicating that they captured intrinsic spike-to-spike variation; only these sessions were used where the intrinsic–extrinsic decomposition was needed (Fig. 6E,F), whereas analyses based on distance and residual magnitude used all 16. Because the predictions could, in principle, miss burst-related waveform change, and a large residual could reflect poor model fit, various validations are presented later that confirm predicted variation is largely burst-related and that the unexplained variation is largely consistent with spike overlaps. Note that drift is a third major presumed source of error, and we return to it below.

**Figure 5:**
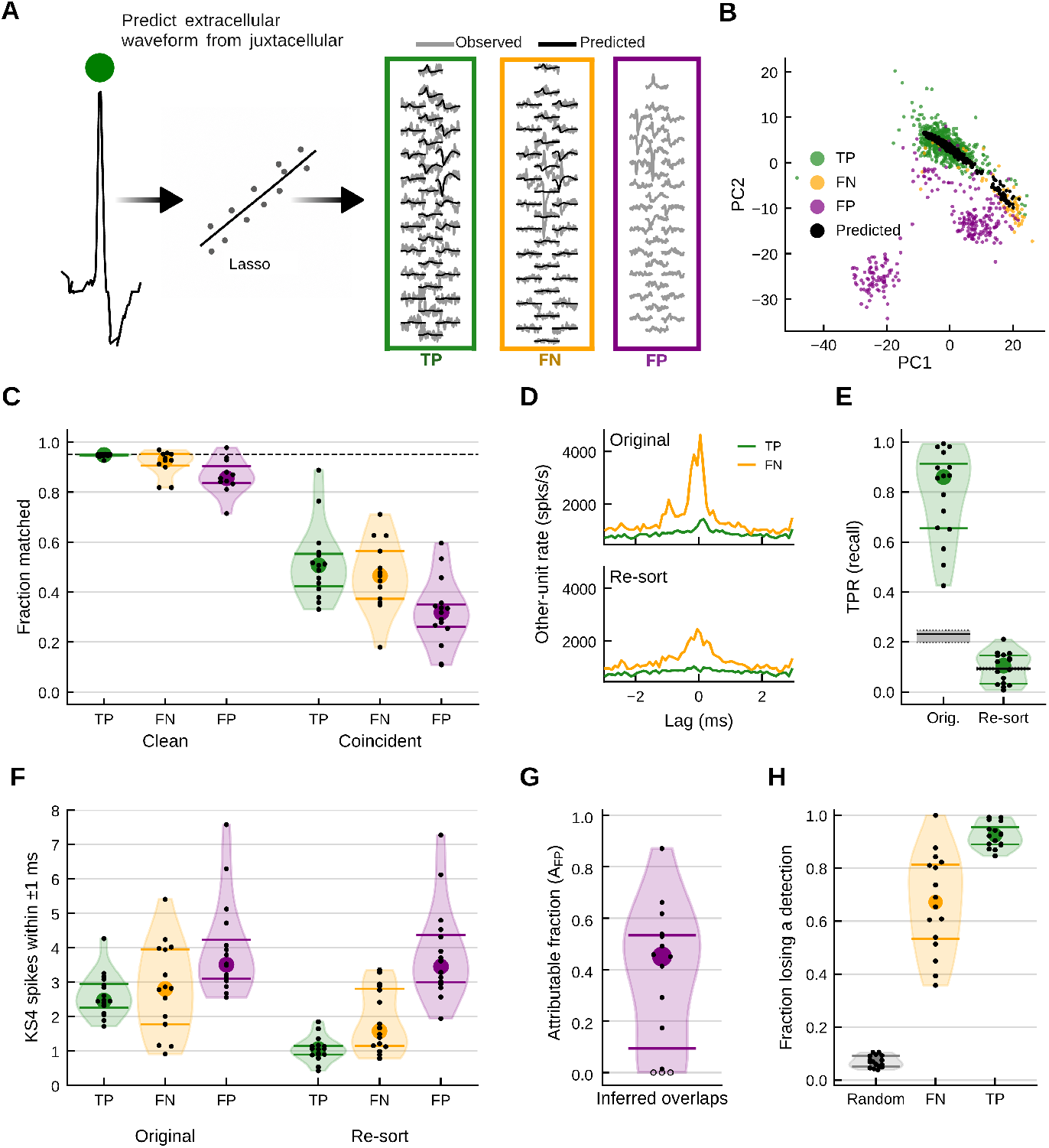
Many false positives arise from wrongly inferred spike overlaps, and many false negatives appear to be detected but assigned to other units. (A) Schematic: the extracellular waveform of each ground-truth spike is predicted from its juxtacellular waveform by LASSO regression; FP waveforms are compared with the *distribution* of predictions. (B) FP, TP and FN waveforms and TP and FN predictions projected onto two principal components in an example neuron. (C) Fraction of TPs, FNs and FPs *matched* for clean and coincident events (clean, no other KS4 unit within ±1 ms); dashed line, 0.95, the expected value for clean ground-truth spikes. Coincident TPs and FNs are weighted to the FPs’ number of nearby spikes. (D) Rate of other units’ detections around TP and FN times in an example neuron (THA-2), in the original sort and after subtracting the ground-truth neuron and re-sorting. (E) Recall of the best-matching unit in the original sort and the re-sort. Line and band, median and 95% interval across shuffles of chance recall, from shuffling each sort’s own unit labels with unit sizes preserved. Note, for the resort the uncertainty band is very thin. (F) Mean number of KS4 spikes within ±1 ms of TPs, FNs and FPs in the original sort and the re-sort, including the event’s own detection where present. (G) Estimated fraction of FPs attributable to inferred overlaps (*A*_FP_); hollow dots, negative estimates shown at zero. (H) Fraction of random times, FNs, and TPs that lost a nearby detection after subtraction and re-sorting. Violins, distribution across neurons; dots, individual neurons. KS4 NM unit, drift-off sort; *n* = 16 neurons (11–15 per group in C; 15 values in G).

In this section, we use the predicted waveform to examine the contribution of putative spike overlap to sorting errors (the next section will study the relative contributions of overlap versus bursting to error). FNs and FPs tended to relate to the predicted extracellular waveforms differently (see an example neuron’s waveform embeddings in Fig. 5B). We call an event *clean* if no other KS4 unit had a spike within ±1 ms, not counting units that were putative oversplit clusters of the ground-truth neuron (units other than the NM unit with precision *>* 0.5 against the ground truth), and *coincident* otherwise. As a baseline for how much the ground-truth neuron’s waveforms vary in the absence of overlaps, for each *clean* ground-truth spike we computed the Euclidean distance between its recorded extracellular waveform and the nearest predicted waveform *other than its own*, in the full space of whitened waveforms. We measured the same distance for every TP, FN, and FP, and called an event *matched* if its distance fell below the 95th percentile of the clean ground-truth distances, cross-fitted so that no clean spike set its own baseline (Methods). So, we now have, for any set of extracellular events, a simple measure of how much their waveforms look like clean ground-truth spikes, likely those without an overlap (as judged by KS4). For both clean and coincident events, FPs were matched less often than TPs and FNs, whereas FNs were matched almost as often as TPs (Fig. 5C). Among clean events, the median fraction matched was 0.95 for TPs (0.95 is the expected value for clean ground-truth spikes by construction), 0.93 for FNs, and 0.86 for FPs (FN vs. FP, *p* = 0.047); among coincident events, with TPs and FNs weighted to the FPs’ number of nearby spikes, it was 0.49, 0.47, and 0.29 (FN vs. FP, *p* = 0.003); TP vs. FN was not significant for either event (exact sign test *p* = 0.55 clean, *p* = 1 coincident). This means, by Euclidean distance, the waveforms KS4 *rejected* (FNs) resembled the ground-truth neuron’s predicted waveforms nearly as closely as those it *accepted* (TPs), and more closely than those it *misattributed* (FPs).

We next wished to compare how often TPs, FNs and FPs coincide with other units’ detections, but this comparison is confounded: a TP’s count always includes its own detection by the NM unit, whereas an FN might occur because it was missed outright, or because the spike was mislabeled. In the latter case, the mislabeled detection would inflate the count of other units’ coincident detections. Excluding putative oversplit clusters here would remove only part of this bias, since a mislabeled spike can be assigned to any unit, not only to potential oversplit clusters of the ground-truth neuron. To correct for this, we artificially removed the ground-truth neuron from each recording and sorted it again (Methods): at every ground-truth spike, the predicted extracellular waveform was subtracted from the recording, and KS4 was rerun with the same settings. We show an example neuron (Fig. 5D), where other units’ detections around FN times peaked sharply at zero lag in the original sort, and this peak was much reduced after re-sorting whereas the rate around TP times changed little, indicating that the excess spikes in the original sort were mislabeled (note, however, that in this example the remainder reveals that many FNs appear to have occurred due to spike overlaps). The recall of the best-matching unit fell from a median of 0.86 (IQR 0.66–0.91) in the original sort, far above a chance value of 0.23 expected from a random clustering of the original sort’s detections (95% interval across shuffles 0.20–0.25), to 0.10 (IQR 0.03–0.15) in the re-sort, also close to the 0.09 expected by chance in the re-sort (0.09– 0.10; Fig. 5E), indicating that the neuron had been artificially removed effectively. As a brief digression, note that this analysis also provides evidence that all ground-truth neurons were indeed analyzed by KS4, complementing our reports of extracellular SNR (Fig. S5B).

So, let us now return to the question of why FP waveforms were matched less often than FN waveforms. Assume there are essentially two *systematic* ways an FP might plausibly occur in template-based sorters:

- A *similar-template FP* occurs when a nearby neuron produces a waveform resembling the ground-truth neuron’s, perhaps with distinctions suppressed by featurization.
- An *inferred-overlap FP* occurs when the sorter infers a spike overlap at a particular time and erroneously attributes residual voltage to the ground-truth unit.

For clarity, note that an inferred-overlap FP necessarily implies coincident detections in KS4’s output (i.e., it refers to the sorter’s belief about overlaps) but a coincident detection does not imply an inferred-overlap FP. If inferred-overlap FPs are present, more KS4 spikes from other units should occur near FPs, on average, than near TPs, where the latter give a baseline coincidence rate for correctly detected spikes. To confirm this, we counted the KS4 detected spikes within ±1 ms of each TP, FN, and FP, including the event’s own detection where present (Fig. 5F; Methods). KS4 reported more spikes near FPs than TPs (median across neurons of the mean number of KS4 spikes within ±1 ms, 3.51 for FPs vs. 2.47 for TPs; higher for FPs in 16 of 16 neurons, two-sided sign test *p* = 3.1 × 10^−5^).

Unlike inferred-overlap FPs, similar-template FPs should not *necessarily* depend on the coincidence rate. If so, similar-template FPs should, on average, have about as many coincident detections as TPs. Therefore, the mean shift and skewed tail for FPs observed in Fig. 5F likely indicate that FPs arise from a *mixture* of similar-template FPs, with a coincidence rate that is in-distribution with TPs, and inferred-overlap FPs, which scale with the coincidence rate (more on this later).

In the original sort, FNs had about as many nearby KS4 spikes as TPs (median 2.80 vs. 2.47; higher for FNs in 6 of 16 neurons, two-sided sign test *p* = 0.45). However, as mentioned before, this comparison is confounded by mislabeled FNs. Assuming perfect subtraction of the ground-truth waveforms, the re-sort removes this problem because no ground-truth spike retains its own detection and thus the reported counts estimate the number of other units’ detections at each time on the same footing for all three event types. In the re-sort, FN times had more nearby detections than TP times (1.58 vs. 1.05; 14 of 16 neurons, two-sided sign test *p* = 0.0042), and FPs had the most coincident detections by an even wider margin (3.45; higher than FNs in 13 of 16 neurons, *p* = 0.021, and higher than TPs in 16 of 16, *p* = 3.1 × 10^−5^; Fig. 5F). FNs thus occurred amid more activity of other neurons than TPs (we will return to this in Fig. 6D), though still less than FPs did. Note also that the FP distribution was less skewed in the re-sort, confirming the earlier premise: the NM unit’s own detection had given every FP a floor of at least one coincident spike, and in the re-sort, while the extracellular voltage was unchanged for FPs, the sorter no longer had the ground-truth neuron’s waveform to explain it with, and often placed no extra detection there at all for many units. Nevertheless, while the distribution was less skewed, the median nearby-spike count at FP times was nearly unchanged in the re-sort, indicating that the sorter often found an alternative explanation for the residual voltage.

**Figure 6:**
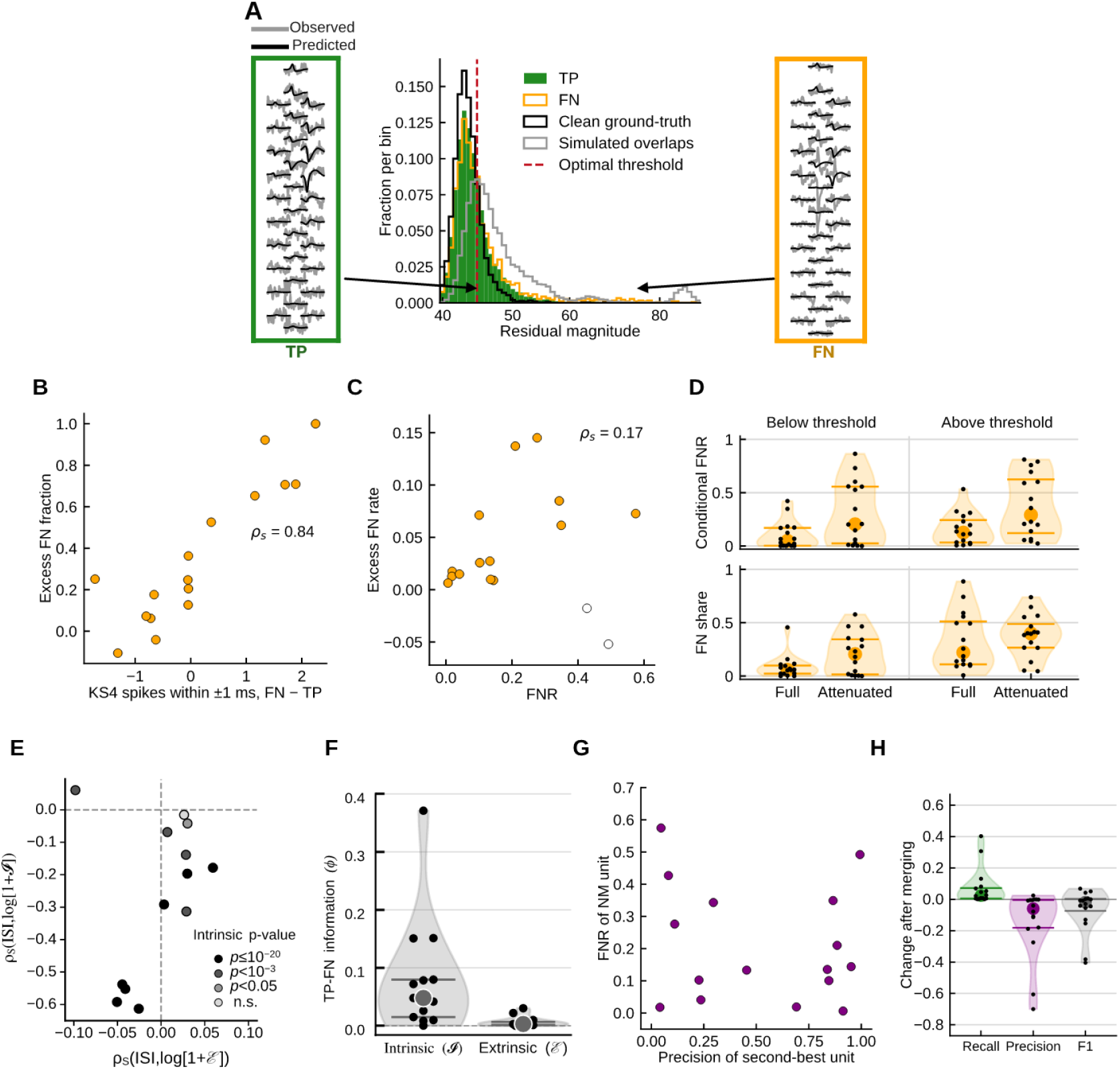
The relative contribution of intrinsic and extrinsic sources to sorting error, and the limits of merging. (A) Residual magnitudes for one neuron’s TPs and FNs, with clean ground-truth spikes and simulated spike overlaps defining the optimal threshold that separates clean spikes (putative non-overlaps) from putative overlaps. (B) Excess FN fraction (FNs above the threshold beyond the below-threshold rate, as a fraction of FNs) versus the per-neuron difference between FNs and TPs in the number of nearby KS4 spikes; the waveform-based and timing-based measures of overlap prevalence agree. (C) Overlap-related FNs per ground-truth spike (excess FN rate) versus FNR; hollow dots, negative estimates (spikes above the threshold had a lower FNR than those below it likely because of noise). (D) Top, conditional FNR of full-height and attenuated spikes (Fig. 3F), below and above the optimal threshold (A). Bottom, each group’s fraction of the neuron’s FNs. (E) Per-neuron Spearman correlation of preceding ISI with extrinsic (x-axis) and intrinsic (y-axis) waveform variation. (F) Held-out TP-FN information (*ϕ*) carried by intrinsic and extrinsic waveform variation. (G) Precision of the next-cleanest unit versus the NM unit’s FNR. (H) Change in recall, precision, and F1 on merging that unit into the NM unit. Violins, distribution across neurons; dots, individual neurons. KS4 NM unit, drift-off sort; *n* = 16 neurons (13 in E and F).

Neither the re-sort nor the counts in Fig. 5F can, by themselves, estimate what fraction of FPs are inferred-overlap FPs. The re-sort measures the fraction of events that lose a detection, and for FPs there is no group analogous to the TPs that we can use to normalize that fraction. Furthermore, while clean FPs should only be similar-template FPs, coincident ones could be either, since similar-template FPs *can also be coincident*. To roughly isolate the two parts of the mixture, we compared error rates between clean and coincident detections. Assuming that KS4 was able to detect when no true overlaps occurred, the FDR among clean detections provides the baseline attributed to similar-template errors. FPs among coincident detections in excess of that rate, as a fraction of all FPs, then estimate the fraction of FPs attributable to inferred overlaps (this reasoning relies on an equal-rate assumption; see Methods). This estimate, *A*_FP_, had a median of 0.45 (IQR 0.09–0.53, *n* = 15 neurons; Fig. 5G). Negative estimates of *A*_FP_, which mean the FDR was higher among clean than coincident detections (Methods), are plotted at zero in the figure but reported here: CA1-6, −0.20; SUB-1, −0.18; THA-4, −2.05, a neuron with 17 FPs. We will return to an analogous analysis of FNs in the next section.

As a byproduct of this analysis, the re-sort also estimated how many FNs KS4 detected but assigned to another unit. A *mislabeled* FN is a detection of a ground-truth spike, so it should disappear when the neuron is removed, as a TP’s detection does, whereas a spike missed outright leaves nothing to remove (i.e., nothing the sorter considered to be statistically meaningful). After subtraction and re-sorting, 92% of TPs lost a nearby detection in the median neuron (IQR 89–96%), compared with 7% of random times (IQR 5–9%), and 67% of FNs (IQR 53–81%; Fig. 5H); FNs lost a detection more often than random times in 16 of 16 neurons (two-sided sign test *p* = 3.1 × 10^−5^), and TPs more often than FNs in 15 of 16 (*p* = 0.00052). Correcting for the TPs’ incomplete detection loss, within attenuated and full-height spikes separately (Methods), the estimated fraction of a neuron’s FNs that were detected but assigned to another unit had a median of 0.77 (IQR 0.57–0.91, *n* = 16). Therefore, the majority of FNs appear not to have been missed outright but were instead mislabeled.

To summarize this subsection, we argued FPs are likely to arise from two systematic sources: similar templates and inferred overlaps. Inferred-overlap FPs seem to account for roughly half of all FPs in the median neuron, and that fraction varied widely across neurons. FNs mostly appear to have been detected but assigned to other units (Fig. 5H), and occurred amid more activity of other neurons than TPs did (Fig. 5F), though less than FPs did. That FPs are especially elevated during spike overlaps is consistent with the previous LFP result for FPs (Fig. 4). These results suggest that, while the overall precision of KS4 was comparatively good, a sizable portion of the remainder might be best managed by handling putative overlaps more skeptically. Put more cautiously, if one’s aim is to boost recall by resolving overlaps, restraint is advised: within the present paradigm, further emphasis on resolving spike overlaps might raise FDR more than it lowers FNR.

#### Bursting and spike overlap

Let us now compare the contribution of spike overlap and bursting more directly. We quantify the contribution of probe drift separately below. Information about bursting is available on the juxtacellular channel for FNs, so we reuse the LASSO predicted extracellular waveforms from the previous section. For each ground-truth waveform, let the *residual magnitude* be the norm of its recorded waveform minus its predicted waveform. In Fig. 6A, for example, the residual magnitude for the illustrated TP is small, whereas it is large for the FN evidently because of a spike overlap.

In the previous section, we used the number of KS4 units detected in ±1 ms as a proxy for spike overlaps. This proxy might have reflected the sorter’s beliefs about spike overlap (hence our earlier term ‘inferred-overlap FP’). Here we attempt to isolate the contribution of *true* overlaps a bit more sharply, although still relying on instances when KS4 judged there to be no other-unit spikes near the ground-truth spike. To approximate a typical residual magnitude consistent with true spike overlaps, denote *clean* ground-truth spikes, in the same sense as before, as those with no spike of another KS4 unit within ±1 ms. We *simulate* spike overlaps by adding the mean waveform of another unit, at a random lag within ±1 ms, to the recorded waveforms of clean ground-truth spikes. Separately for each neuron, we calculated the residual magnitude that best separated clean ground-truth spikes from simulated overlaps, denoted the *optimal threshold* (Fig. 6A). The optimal threshold separated clean from simulated overlaps with a median AUC of 0.78 (IQR 0.77–0.82; median sensitivity 0.70, specificity 0.76).

As a validation, we checked that this waveform-based measure of overlap agrees with the timing-based one of the previous section. We define the *excess FN fraction* as the number of FNs above the threshold in excess of the number expected at the below-threshold FNR, as a fraction of all FNs (Methods). Across neurons, the excess of nearby KS4 spikes at FNs relative to TPs (Fig. 5F, original sort) closely tracked the excess FN fraction (*ρ_s_* = 0.84, *p* = 5 × 10^−5^; Fig. 6B). Overlaps inferred from KS4’s spike timing and overlaps estimated via large waveform residuals thus approximately coincide, which supports both the timing-based analyses above and the use of the residuals as a measure of extrinsic waveform variation in what follows.

But how many FNs might *true spike overlaps* account for? Many FNs participating in spike overlaps would presumably still have been missed because of bursting had the overlap not occurred, which is why the excess FN fraction defined above compares spikes above the threshold with the below-threshold FNR; it is thus a distinct estimate of the fraction of FNs attributable to spike overlaps. The median neuron had 25% of its FNs attributable to spike overlaps by this measure (IQR 11–67%, *n* = 16; Fig. 6B); in two neurons the estimates were negative (NEO-2, −0.11; SUB-2, −0.04), meaning their above threshold spikes were missed less often than their below threshold spikes, contrary to the assumption that overlaps add misses on top of the below-threshold rate (we retain calling the metric a fraction for when the assumption holds, and still plot the exceptions).

We next wondered whether FNs attributable to spike overlaps can explain the wide variability in FNR across neurons observed earlier (Fig. 2D). Suppose some neurons had lower recall than others only because they fire during ripples, when population activity is elevated. Then the neurons with the highest FNR would also be the ones with spikes that are most often overlaps, and the number of overlap-related FNs per ground-truth spike (the *excess FN rate*) would rise with FNR. This picture appears too simplistic: overlap-related FNs were about 2% of ground-truth spikes in the median neuron (IQR 1–7%) and were not significantly related to FNR across neurons (*ρ_s_* = 0.17, *p* = 0.52, *n* = 16; Fig. 6C). Two neurons had small negative estimates (NEO-2 and SUB-2; hollow dots), consistent with overlaps adding few FNs there, although their thresholds separated overlaps as well as in other neurons (AUC 0.77 and 0.85, respectively). Excluding these two neurons from the analysis, the excess FN rate was moderately positively associated with FNR (*ρ_s_* = 0.65, *p* = 0.011, *n* = 14). Even so, the fraction of FNs attributable to overlaps fell sharply as FNR rose (*ρ_s_* = −0.77, *p* = 0.0005). Thus, *considered on their own*, overlaps were associated with only a modest excess of FNs in most neurons but accounted for little of the variability in FNR across neurons, contrary to what the ripple thought experiment above imagines. However, overlaps might *interact* with burst attenuation, which we consider next.

#### Distribution of FNs

To improve spike sorting, it is useful to quantify how difficult a particular kind of event is to detect (its conditional FNR) and how much of the total loss that kind of event accounts for (its FN share), which weighs its difficulty by how commo it is. Therefore, we returned to our earlier analysis of burst-attenuated spikes, as measured juxtacellularly, and split each neuron’s spikes by attenuation (Fig. 3F) and by whether their residual magnitudes fell below or above the optimal threshold (Fig. 6D; Methods). Full-height spikes below the threshold were rarely missed (median conditional FNR 0.05, IQR 0.00–0.17), and attenuated spikes below the threshold had a median FNR of 0.20 (IQR 0.02– 0.56); above the threshold, the median FNR of full-height spikes was 0.13 (IQR 0.03–0.24) and the median FNR of attenuated spikes was 0.29 (IQR 0.12–0.62). Both burst-attenuation and overlap-like residuals raised the FNR in most neurons, whether the other factor was present or not (14 neurons in each of the four comparisons, *p* = 0.0042, two-sided sign tests). We also computed, within each neuron, how much each factor raised the FNR relative to full-height spikes below the threshold. Across neurons, the median increase was 2.5-fold for burst-attenuation alone (IQR 1.8–8.5), 2.2-fold for overlap-like residuals alone (IQR 1.4–4.1), and 7.2-fold for the two together (IQR 2.7–23.8). Two neurons with reference FNRs exactly at zero were excluded (not plotted). Burst-attenuated spikes with overlap-like residuals were also the only group over-represented among FNs, making up 22% of a neuron’s spikes (IQR 11–26%) but 40% of its FNs (IQR 27–49%), which held in 15 of 16 neurons (*p* = 0.00052; Fig. 6D, bottom). In the other three groups, the share of FNs was about the same as, or far below, the share of spikes: 29% of spikes and 22% of FNs for full-height spikes above the threshold, 18% and 21% for attenuated spikes below it, and 26% and 6% for full-height spikes below it.

We also ran this analysis with spikes split by whether another KS4 unit was detected within ±1 ms, as in Figure 5 (not plotted). As before with the re-sort analysis (Fig. 5H), we defined coincidence after the ground-truth neuron had been subtracted. This gave a highly similar quantification: conditional FNR 0.03, 0.21, 0.13 and 0.30 for full-height clean, attenuated clean, full-height coincident, and attenuated coincident spikes. From these analyses, since spikes that are both attenuated and overlapped are over-represented among FNs, we suggest that next-generation sorters may try to address both problems at once.

#### Probe drift and slow nonstationarity

Spikes can also be missed when the neuron moves relative to the probe, or when the recording changes in other ways over time, such as the cell’s health declining. We therefore asked how much of the error varied over the course of a recording. Binning each session into ten equal intervals, recall varied monotonically with time in 3 of 16 sessions after Bonferroni correction (8 of 16 uncorrected; exact permutation test on the Spearman correlation). Enabling KS4’s drift correction left median recall almost unchanged (NM, 0.855 vs. 0.847 with it off; Table S2). With correction enabled, the Spearman correlation between recall and time fell below significance in three sessions and decreased in magnitude in two, but rose in magnitude in five others. Drift correction therefore helped some recordings and hurt others, with little net effect on recall.

To ask whether this time-varying error also shaped the distribution of FNs across the four groups of Fig. 6D, we removed, within each neuron, the 30% of time bins with the lowest recall, which lowered the median FNR from 0.14 to 0.10, and recomputed the split on what remained. Every comparison was preserved or increased: the conditional FNRs fell proportionately (0.03, 0.14, 0.08 and 0.24 for the four groups), burst-attenuated spikes with overlap-like residuals remained the only group over-represented among FNs (16% of spikes, 37% of FNs, in 16 of 16 neurons, *p* = 3.1 × 10^−5^), and the per-neuron fold changes were unchanged or larger (3.0-fold for attenuation alone, 2.8-fold for overlap-like residuals alone; not plotted). We did not pursue drift nonstationarity as a third factor in this split, since we lack the kind of independent measurement for it that the juxtacellular recording gives us for bursting and the waveform residual gives us for overlap. To conclude, although slow time-varying error was prominent in a few recordings, it did not explain the relative ordering or qualitative contrasts among the four groups’ FNRs.

#### The most predictive errors

Not that the share of FNs a particular situation accounts for post hoc is not the same as an error that can be anticipated, and it is thus also useful to quantify how *predictable* a particular kind of error is. Therefore, we next wondered which kind of waveform variation predicts which spikes are FNs in held-out data. To compare the contributions of intrinsic (e.g., bursts) and extrinsic (e.g., spike overlap) factors, we calculated two measures for each ground-truth waveform. The first, intrinsic variation, is defined via the deviation of the ground-truth waveform’s predicted waveform from those of spikes preceded by at least 200 ms of silence (presumably not burst-affected). Note that, here we apply the analysis to the extracellular, rather than juxtacellular, waveforms in contrast with the previous burst-attenuation analysis. The second measure, extrinsic variation, is defined via the residual between the observed ground-truth waveform and its predicted waveform (Methods). Among the 13 neurons meeting the waveform-prediction criterion, intrinsic variation increased at short preceding ISIs (*ρ_s_ <* 0 in 12 of 13 neurons; median *ρ_s_* = −0.20; Fig. 6E), confirming again that the predicted waveform changes are largely burst-related. In contrast, correlations between ISIs and extrinsic variation *ε* were small and scattered around zero (median *ρ_s_* = 0.01, range −0.098 to 0.059; |*ρ_s_*| ≤ 0.10 in all 13 neurons). Intrinsic variation also carried more held-out information about which spikes were FNs than extrinsic variation did (11 of 13 neurons; median *ϕ_I_* − *ϕ_ε_* = 0.045, 95% interval across surrogates 0.0436–0.0467; one-sided exact paired sign-flip test, p=0.011). The intrinsic measure depends only on the neuron’s own predicted waveform, whereas the residual includes spike overlaps on distant channels, which likely do not affect detection.

#### Merging

We next asked whether false negatives could be recovered by merging. Merging is usually guided by refractory CCGs, but KS4 already merges clusters whose CCGs are refractory and whose templates are similar, so few such candidates remained: only 3 of 16 neurons had any cluster with a refractory cross-correlogram against the NM unit, and those clusters contained almost no ground-truth spikes (not plotted). We therefore chose as the candidate the cluster with the highest precision against the ground-truth neuron other than the NM unit. This choice departs slightly from the BM procedure of Fig. 2, perhaps better matching how a human curator would merge units. In the median neuron, the precision of the candidate cluster was 0.57, but it varied widely (IQR 0.20–0.87) and was unrelated to the neuron’s FNR (*ρ_s_* = −0.01; Fig. 6G). Merging it raised recall by a median of only 0.02 (IQR 0.01–0.07) while lowering precision by 0.06 (IQR 0.00–0.18), so F1 fell in 11 of 16 neurons and rose by more than 0.05 in only two (Fig. 6H). In five neurons, merging did not lower precision, but the clusters were small and added little recall; the two neurons in which F1 increased by more than 0.05 were among these. In three others, the cluster’s precision was high but its spikes fell almost entirely at ground-truth spikes the NM unit had already detected, so merging added almost no recall. In the remaining eight, even the highest-precision cluster had low precision (full range across neurons 0.04–0.45), so merging lowered precision overall. Even the best available merge therefore cost more in precision than it gained in recall, which is why neither the earlier BM procedure of Figure 2 nor the single merge here could recover the missing spikes in most neurons.

### Testing across spike sorters

To ask whether the observed error patterns thus far reflect KS4 specifically or something more general, we evaluated three additional spike sorters representing distinct algorithmic approaches: IC, MS5 and SC2. KS4 combines graph-based clustering with template deconvolution; IC uses density-peak clustering and waveform-based cluster merging without template deconvolution; MS5 uses ISO-SPLIT clustering followed by classifier-based spike assignment; and SC2 uses matched-filter detection, density-based clustering, and template matching. We selected these from among the best-performing sorter families in the KS4 benchmark (Pa-chitariu et al., 2024). Note that MS5 and SC2 required parameter adjustment while KS4 and IC were run at their defaults, so differences across sorters could partly reflect parameter configurations rather than the algorithms. MS5 in particular under-clustered severely on a probe of this geometry with its default settings, so we adjusted three of its parameters (Methods).

#### Overall performance comparison across sorters

Under NM, median F1 ranked KS4 (0.849) above IC (0.813), MS5 (0.732) and SC2 (0.644), and the same ordering held under BM (Fig. 7A; Table 2). KS4 had higher F1 than MS5 and SC2 under both merge conditions (*p* ≤ 0.025), and IC had higher F1 than SC2 under NM (*p* = 0.039); no other pair differed after correction, including KS4 and IC. Notably, IC had the highest precision of the four sorters, and was the only one whose F1 was unchanged by BM. BM merged more than one unit in 9 of 16 sessions for SC2, more than twice as often as for any other sorter. This recovered much of its recall (0.685 to 0.895) at a substantial cost in precision (0.825 to 0.741), so its F1 improved only modestly, suggesting the merged units were largely not the ground-truth neuron’s. Variability across sessions was greater in recall than in precision for KS4, MS5 and SC2, as in Figure 2, but not for IC.

**Figure 7:**
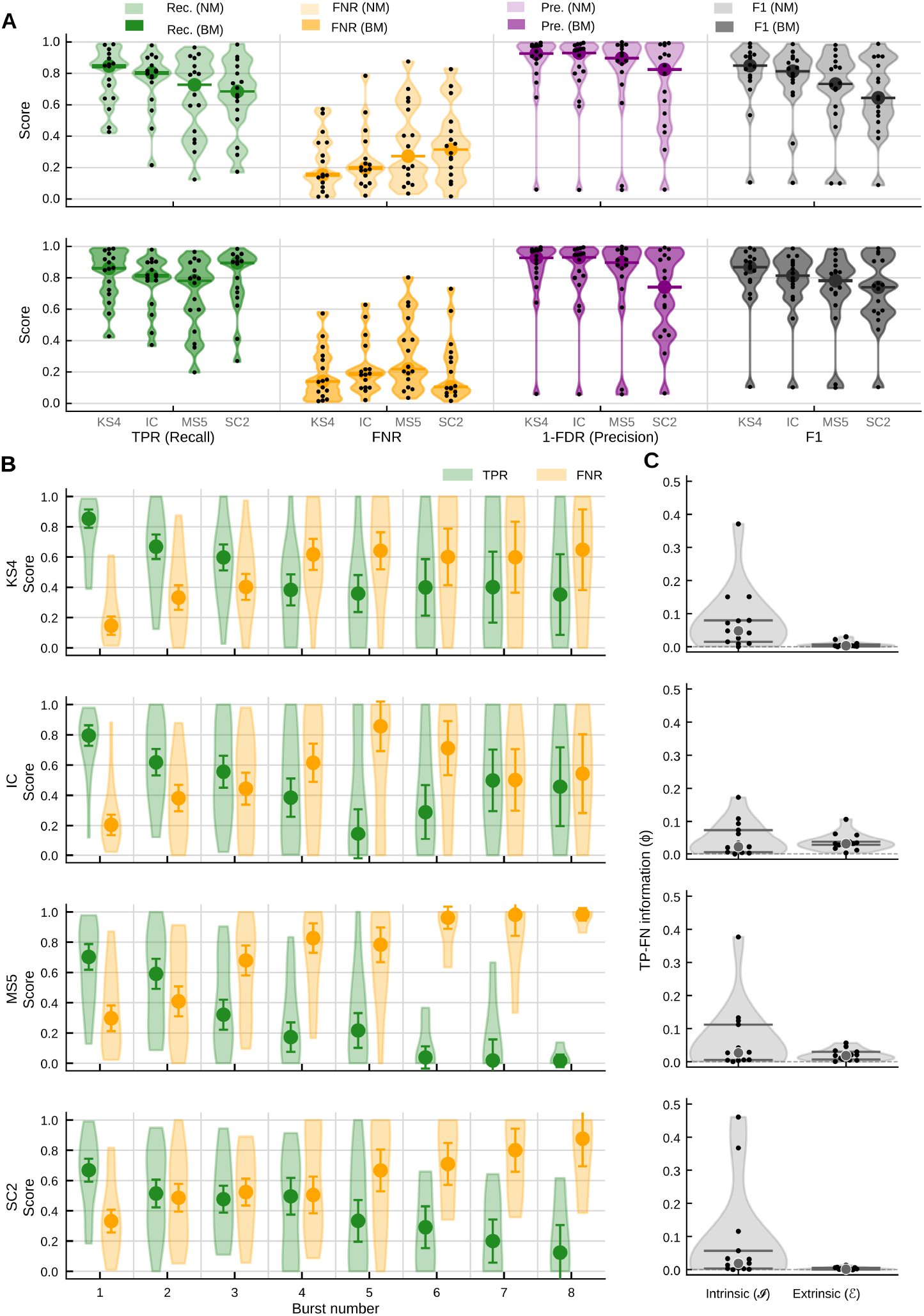
Testing of main effects across spike sorters. (A) Overall performance of KS4, IC, MS5, and SC2 under no merge (NM) and best merge (BM); one dot per session. (B) Conditional recall and FNR versus burst position under NM, one row per sorter. Violins show distributions across sessions; dots indicate medians and bars indicate ±1 SEM. The number of contributing sessions decreases from 16 at burst position 1 to 4 at position 8. (C) Information carried by intrinsic waveform variation, *I*, and extrinsic residual variation, *ε*, about whether ground-truth spikes were missed by KS4, IC, MS5, and SC2. Dots individual neurons, horizontal bars mark Q1 and Q3. Sorters are ordered by median NM F1 throughout; *n* = 16 neurons in A and B, 13 in C (those with informative cross-fitted waveform predictions).

**Table 2:** Spike sorting performance in four sorters across 16 sessions. Values are medians across sessions (interquartile range); 95% intervals across surrogates were within 0.01 of each median and are omitted. KS4, MS5, and SC2 were run with drift correction off; IC was run at its default settings, which include its built-in drift handling.

| Merge | Metric | KS4 | IC | MS5 | SC2 |
|---|---|---|---|---|---|
| **NM** | Recall | 0.847 (0.642–0.903) | 0.804 (0.715–0.863) | 0.727 (0.418–0.859) | 0.685 (0.553–0.811) |
|  | FNR | 0.153 (0.097–0.358) | 0.196 (0.137–0.285) | 0.273 (0.141–0.582) | 0.315 (0.189–0.447) |
|  | Precision | 0.927 (0.846–0.976) | 0.930 (0.788–0.963) | 0.897 (0.789–0.974) | 0.825 (0.525–0.937) |
|  | F1 | 0.849 (0.768–0.912) | 0.813 (0.725–0.885) | 0.732 (0.577–0.851) | 0.644 (0.521–0.863) |
| **BM** | Recall | 0.860 (0.717–0.932) | 0.814 (0.745–0.899) | 0.781 (0.560–0.859) | 0.895 (0.699–0.928) |
|  | FNR | 0.140 (0.068–0.283) | 0.186 (0.101–0.255) | 0.219 (0.141–0.440) | 0.105 (0.072–0.301) |
|  | Precision | 0.927 (0.825–0.976) | 0.930 (0.788–0.961) | 0.897 (0.789–0.973) | 0.741 (0.457–0.928) |
|  | F1 | 0.867 (0.783–0.923) | 0.813 (0.725–0.899) | 0.782 (0.677–0.851) | 0.740 (0.584–0.907) |

#### Burst-related false negatives replicate across sorters

FNR increased with burst number in all four sorters (Fig. 7B). Averaged across sessions, it rose from 22% at the first burst position to 57% at the eighth for KS4, from 25% to 52% for IC, from 35% to 96% for MS5, and from 35% to 78% for SC2. Per-session Spearman correlations between FNR and burst number were positive in 12 of 15 KS4 sessions, 13 of 15 MS5 sessions, and 12 of 15 SC2 sessions (median *ρ_s_* = 0.78, 0.70, and 0.92; two-sided sign test *p* = 0.035, 0.0018, and 0.035). In IC the dependence was evident but non-monotonic, so the rank correlation does not capture it (positive in 8 of the 13 sessions with a nonzero correlation, median *ρ_s_* = 0.50, *p* = 0.58); we did not attempt to model the shape.

#### Intrinsic waveform information tested across sorters

Intrinsic variation carried more held-out TP–FN information than extrinsic variation in 11 of 13 sessions for KS4, 8 of 13 for IC, 9 of 13 for MS5, and 10 of 13 for SC2 (Fig. 7C). Median *ϕ_I_* − *ϕ_ε_* was 0.045 for KS4, 0.010 for IC, 0.008 for MS5, and 0.018 for SC2. The difference survived Holm correction for KS4 and SC2 (one-sided exact paired sign-flip tests, *p* = 0.011 and 0.012), but not for IC (*p* = 0.29) or MS5 (*p* = 0.11).

## Discussion

Spike sorting dates to the early work of Gerstein and Clark (1964), and our subsequent account of population coding (particularly temporal and sequence coding), network dynamics, and functional connectivity has since remained susceptible to structured biases induced by its errors (Pazienti and Grün, 2006; Cohen and Kohn, 2011; Siegle et al., 2021). Using awake juxtacellular ground truth, we found that four current spike sorters perform worse than conventional benchmarks might suggest, and that their errors are structured rather than random. Below we consider what these results mean for the interpretation of existing extracellular datasets and for the development of better spike sorting.

### Bursting

The difficulty bursting presents for spike sorting is widely recognized (Fee et al., 1996; Lewicki, 1998; Harris et al., 2000; Koukuntla et al., 2025), but perhaps still insufficiently emphasized. Small spikes during bursts cannot be dismissed as functionally negligible: although strongly attenuated at the soma, evidence indicates they recover to full amplitude in the distal axon and trigger transmitter release comparable to regular spiking (Apostolides et al., 2016). Bursts have also been proposed to carry information distinct from isolated spikes and to gate synaptic plasticity (Izhikevich et al., 2003; Payeur et al., 2021). One might believe bursting neurons constitute a small proportion of cells in neuronal populations, and while hunting for neurons to record juxtacellularly, experimentalists are perhaps biased to notice bursting neurons, inflating estimates of their prevalence. However, there are nonetheless at least four errors in reasoning committed if burst-induced error is dismissed too readily.

#### Selection bias

The true rate of bursting cells in neuronal populations is unknown. Establishing it would require intracellular or juxtacellular recording in vivo, which yields very low sample sizes. Most single-cell recordings are made in anesthetized animals, yet burst firing is strongly state-dependent, so its prevalence under anesthesia need not reflect the awake brain (Steriade et al., 2001; Sherman, 2001; Steriade, 2004).

#### False assumption of independence of errors

Detection errors across neurons are not generally independent. For example, one neuron’s false negatives, if detected at all, are the false positives of other neurons. Bursting neurons also complicate the detection of overlapping waveforms during template deconvolution, distorting the detection and featurization of neurons other than the bursting neuron in question. This potentially cascades errors across every unit contributing to the same overlapping events, since a poorly explained spike alters the residual against which the remaining templates are matched. Finally, if a bursting neuron is oversplit and the templates of the oversplit units each have *poor precision* and are used for template deconvolution, errors might yet again cascade across the population in hard-to-predict ways. Indeed, here about three-quarters of false negatives appear to have been detected but assigned to other units (Fig. 5H), and the cleanest cluster available to merge had a median precision across neurons of 0.57, so these spikes were mostly absorbed into other units’ clusters rather than cleanly oversplit (Fig. 6G,H).

#### Fallacy of composition in featurization

Bursting neurons exhibit extreme true within-class variability and thus strain variance-maximizing objectives such as PCA, since preserving intra-class variance degrades discriminative directions under finite representational capacity. That is, it is possible that including a bursting neuron A when estimating a PCA basis could cause two distinct neurons B and C to merge into one cluster rather than two, because the basis is estimated from the covariance of a population of unlabeled spikes.

#### Base rate fallacy

Even if bursting neurons are rare, the relevant quantity is not the proportion of neurons that burst but the bias bursting introduces into reported effects and, by the preceding points, such effects need not have anything to do with bursting. A large share of the scientific questions asked with multiunit recordings concern weak but regular temporal structure among parallel spike trains: multineuronal sequences, functional connectivity inferred from spike timing, responses measured relative to stimulus onset, low-rank population dynamics. Temporally-structured burst events incorrectly assigned to other units are likely to masquerade as inter-neuronal relationships, biasing a much higher proportion of the effects of interest than the proportion of neurons that burst.

That the bursting result holds across KS4, IC, MS5, and SC2 (Fig. 7B) bears on how these errors should be understood. Four sorters with different featurizations, clustering algorithms, and deconvolution strategies fail similarly as a function of burst position, which might indicate that a new paradigm is needed rather than post hoc merging.

### Discriminative power and invariance

Mospike sorters are heuristic pipelines of many independent steps, and an error induced by an early step such as whitening or detection may not manifest until featurization. Nonetheless, a unifying way to think about spike sorting is to conceptualize all steps prior to and including featurization as representation learning. On that view, the operative assumptions are invariance assumptions in this process of representation: if two dissimilar raw waveforms map to similar embeddings, the representation has committed to an invariance. Several are explicit enough to name in KS4, for example. Time-translation and noise invariance are sought through template matching, collision invariance through spatially local dimensionality reduction and deconvolution, probe-movement invariance through drift correction, and a weak form of burst invariance through per-spike rescaling coefficients during deconvolution.

Discriminative power and invariance trade against one another, a trade-off studied in computer vision and machine learning more broadly (Varma and Ray, 2007). An invariance assumption enlarges a statistical model’s acceptance region, which may improve recall but also assuredly degrades precision. The performance patterns reported here are qualitatively consistent with what this trade-off predicts. The strongest invariance assumptions are made to recover spikes from spike overlaps: template deconvolution explains the recording as a sum of templates, and so assumes that each neuron’s waveform has a characteristic size and shape. Characteristically, false positives are then elevated when this assumption breaks: a substantial fraction of false positives occurred during putative overlaps, and false positive waveforms resembled the neuron’s predicted waveforms less often than false negatives did (Fig. 5C,F,G; Fig. 4C; Fig. S5). Drift correction is another such invariance assumption, and enabling it likewise elevated false positives slightly (Fig. S5D). In contrast, KS4 allows only weak scale invariance to accommodate burst-like amplitude variation, but bursts produce substantial changes in the extracellular waveform that grow at short interspike intervals (Fig. 6E), and recall conditioned on burst position was accordingly poor (Fig. 3C). Errors were over-represented where bursting and spike overlap co-occurred, exactly where precision and recall must be traded most delicately, and where uncertainty quantification would therefore help most (see the final paragraph).

These errors were not particular to KS4. IC, MS5 and SC2 differ from it at every step of the pipeline (detection, featurization, and clustering), yet recall degraded with bursting in all four (Fig. 7B). It was not tractable to vary the assumptions of these sorters in a controlled manner, so this analysis did not isolate which assumptions are responsible. However, four algorithms built on different principles failing in similar ways seem to reveal something more general, and makes taking bursting seriously as a design problem worthwhile.

How *predictable* these errors were, though, differed across sorters. While burst-attenuated spikes with overlap-like residuals were the largest share of FNs in KS4, intrinsic waveform variation contributed more held-out information about which spikes were FNs than extrinsic variation did, significantly so in KS4 and SC2 but not in IC or MS5 (Fig. 7C). The two sorters in which intrinsic variation was significantly more informative, KS4 and SC2, both match templates to the recording, whereas IC does not use template deconvolution and MS5 uses classifier-based spike assignment. This is consistent with the idea that a sorter committed to a fixed template is perhaps more robust to collisions but also more susceptible to errors from burst-related changes in waveform. Importantly, burst-related increases in FNR were not limited to template-matching-based sorters: IC and MS5 also varied strongly with burst number. Thus, the effect of bursting was general, while the relative predictive value of intrinsic and extrinsic variation differed across sorters. Where the effect was present, we presume it arose because burst-related attenuation produces errors that exhibit a statistical regularity, whereas whether an overlap produces an error likely depends on the fine details of how the overlap occurs, such as its location on the probe, its timing relative to the spike, and the amplitude of the overlapping unit. While detecting that regularity here required ground truth, the statistical regularity itself does not, and might serve as an effective prior for inference in both types of sorters.

### Soft ground truth

“Ground truth” in practice refers to a signal measured at much higher SNR than the method being benchmarked, not to a signal known with perfect confidence. The term was popularized in remote sensing, where surface measurements were used to calibrate satellite imagery (Hoffer, 1972), and that field has since questioned whether the word is appropriate, given that such measurements carry their own error (Woodhouse, 2021). Machine learning treats ‘label noise’ as a serious practical obstacle for supervised learning, with a substantial literature on robust learning under imperfect labels (Frénay and Verleysen, 2014). Yet, in neuroscience, ground-truth data are typically hard-labeled toward the experimentalist’s best judgment.

Even in high-SNR neural recordings, the small events late in a burst can be continuous with the noise distribution on the juxtacellular channel (Fig. 1C). If ambiguity of this kind is present in a juxtacellular recording with a SNR far exceeding anything available extracellularly (Table S1), the failure of most contemporary spike sorters to output soft labels is truly indefensible (cf. Lee et al. (2017)). We hope the way soft labels are propagated into downstream analyses here suggests how the same could be done with extracellular soft labels.

The noise model we used for soft labeling is deliberately conservative, so the reported soft TPR is an upper bound, while the FDR is arguably less affected. The reasoning is as follows. Assume juxtacellular detection is easier than extracellular detection in expectation (Fig. 1B; juxtacellular SNR exceeded extracellular SNR in every session, Table S1, and few candidate events had intermediate posteriors, Fig. S2). An event that cannot be confidently judged a spike on the juxtacellular channel is one the sorter is unlikely to have recovered, and the conservative noise model downweights it, reducing the penalty on the sorter in a graded manner. Reported recall is therefore a ceiling. Precision is arguably much less affected by the proposed conservatism: the sorter’s false positives are likely to arise from a much larger set of confounds unrelated to phenomena that influence the juxtacellular channel (e.g., spikes from other neurons with similar templates).

The above reasoning is largely supported in Fig. S3A, which compares KS4 performance under our conservative soft labels with performance under two alternative hard labelings of the ground truth: KS4 itself applied to the juxtacellular channel, and a 5*σ* amplitude threshold (Quiroga et al., 2004). Recall is considerably higher under our labeling (by 0.14 and 0.32, respectively, under NM), suggesting that our concerns about bursting are, if anything, understated: labelings that count more of the ambiguous small events as spikes penalize the sorter more, not less. Precision, by contrast, was nearly unchanged per session: switching labelings shifted a given neuron’s precision by a median of at most 0.010 (per-neuron paired difference across labelings, not the surrogate interval within our labeling), supporting the intuition that the F1 score was not unfairly punished in aggregate because the spike sorter detected many spikes that were not detected juxtacellularly. Note that, as the juxtacellular labeling becomes more permissive and more true noise events are counted as spikes, precision cannot decrease, and in the limit of labeling every event a spike it must reach 1. Recall, meanwhile, can only fall. As we said earlier, our conservative labeling thus roughly gives an upper bound on recall and a lower bound on precision.

### Scope of the present study

This study should not be interpreted as a calibrated prediction of sorting performance as it may be employed generally. Juxtacellular recordings are unstable in awake animals, and these sessions are consequently noisier and shorter than typical neural recordings in other laboratories.

While almost all existing ground-truth data come from anesthetized or in vitro preparations, we are aware of two exceptions. First, a pair of recordings in Allen et al. (2018), in which whole-cell access was obtained under isoflurane anesthesia, which was then withdrawn, yielding ∼7.6 minutes of paired recording from each of two layer 5 neurons in freshly awake, head-fixed mice: roughly fifteen minutes of awake ground truth in total. Second, some of the sessions we recorded here contributed to the SpikeForest paper (Magland et al., 2020), although it appears this paper had quite strict exclusion criteria (see more below). There is an unavoidable tradeoff between validity and certainty, and the scarcity of awake ground-truth data reflects that. That is, the recording conditions most valid for benchmarking (in behaving animals) are exactly those that are most unstable, and so we instead addressed this tradeoff by quantifying uncertainty in the (soft) ground truth itself.

Some false negatives, particularly the smallest late in bursts, may be only weakly present on the extracellular probe, and no sorter could be expected to recover them. However, several observations suggest this does not account for most burst-related error. First, burst-attenuated spikes *without* overlap-like residuals were often detected (median recall 0.80; Fig. 6D), and FN waveforms were nearly as close to the neuron’s predicted extracellular waveforms as typical true positives (by Euclidean distance; Fig. 5C). Second, when the neuron was subtracted from the recording and the data re-sorted, 67% of FNs lost a nearby detection, compared with 92% of TPs and 7% of random times. Thus, correcting for the TPs’ incomplete loss, an estimated 77% of FNs had been detected by KS4 but assigned to other units (Fig. 5H). Third, sorters differed substantially at late burst positions (FNR at the eighth burst position of 57% for KS4 versus 96% for MS5, though fewer sessions contributed there; Fig. 7B), so many such spikes were recoverable by at least one algorithm. In subiculum, the best merge (BM) condition restored much of the missing recall (TPR 0.64 to 0.92; Supplementary Text), although across neurons merging the next-cleanest unit recovered little (Fig. 6G,H). Note also that the conservative soft labels already downweight the smallest, most ambiguous juxtacellular events (Soft ground truth), which are those least likely to be visible extracellularly. Similarly, some spike overlaps may be fundamentally non-identifiable even for an optimal detector, though they receive greater emphasis in the literature than bursting does. Ultimately, however, how blame is divided between probe, preparation, and algorithm arguably matters less than it may seem. What matters most to the scientists who use spike sorters is how much extracellular recording and spike sorting, when taken together, can reveal about how the brain works.

Although the preceding discussion suggests that our data are not strictly comparable to other published benchmarks, it is still reasonable to contextualize how our results compare. SpikeForest matched each ground-truth unit to its best sorted unit within a 1 ms tolerance, averaged over units with SNR ≥ 8, and reported accuracy as *n*_TP_*/*(*n*_TP_+*n*_FN_+*n*_FP_) (Magland et al., 2020). Expressed as *accuracy*, KS4’s median across neurons here was 0.74 (IQR 0.62– 0.84), whereas SpikeForest reported 0.92 and 0.88 for Kilosort and Kilosort2 on the earlier release of recordings from this preparation. The KS4 benchmark instead matched within 0.2 ms, scored each unit as 1 − FNR − FDR, and counted a unit as correctly identified if its score exceeded 0.8 (Pachitariu et al., 2024); in our data, the median score was 0.73 (IQR 0.58–0.83) and 7 of 16 neurons (44%) exceeded 0.8, whereas in their no-drift simulation 92% of simulated units did. Notably, across its paired datasets, nearly all from anesthetized preparations, SpikeForest found precision generally lower than recall (Magland et al., 2020); we see the opposite, consistent with the potential state dependence of bursting.

Two potential concerns arise when comparing our recordings with these benchmarks. First, SpikeForest’s SNR cutoff would have excluded 15 of our 16 neurons. Yet we observed no detectable relationship between performance and extracellular SNR (Fig. S5B); the recall of KS4’s best-matching unit was far above the chance level obtained by shuffling its unit labels in all 16 sessions (Fig. 5E); and subtracting the predicted ground-truth waveforms and re-sorting reduced recall to very near chance levels (Fig. 5E). In the SpikeInterface paper’s simulated recordings, moreover, Kilosort2 achieved near-perfect accuracy for units near SNR 2 (Buccino et al., 2020, Figure 2—figure supplement 1B), and our neurons had a median extracellular SNR roughly twice that (4.1). Therefore, low SNR did not prevent KS4 from attempting to sort these neurons, and their signal quality is thus representative of neurons that neuroscientists might record in other laboratories. The gap in performance relative to these benchmarks is also unlikely due to stricter labeling. As detailed above, we found hard labelings of these data yielded lower recall than our soft labels, not higher (Fig. S3A). Finally, we observed no trend of performance with recording duration (Fig. S3) or with the number of ground-truth spikes per neuron (F1 versus spike count, *ρ_s_* = 0.31, *p* = 0.25; not plotted), which suggests these recordings are long enough to be informative.

Several of our analyses hint at attributing errors to a cause. For example, we defined inferred-overlap FPs only as FPs in excess of an inferred similar-template FP rate. One could instead seek such attributions by modifying the sorter, such as disabling its template deconvolution. But an arbitrarily ablated sorter is not one any developer would recommend and would require intimate knowledge of the subtleties of others’ algorithms. Hence, it was only within the scope of this study to make observational inferences about attribution. Ultimately, the utility of these inferences ought to be tested by whether they prove useful for building better sorters, which will depend on the expertise of the skilled engineers embattled with this difficult problem.

### Ill-posedness and uncertainty

Spike sorting is likely an ill-posed problem, in which case no amount of computation will yield perfect accuracy. Sorters fail in specific conditions— during bursts, during spike overlaps—and those conditions are often the ones scientists care about most. So the assumptions a sorter makes matter whether you build sorters or only use them. Since our results suggest that sorting errors can concentrate in events of scientific interest, that uncertainty should be propagated into the particular effect under study. Furthermore, since an invariance assumption trades false positives against false negatives, the right balance depends on what the recording will be used for. Next-generation sorters might therefore make that balance a function of the scientific question. Single-trial and single-neuron uncertainty quantification is largely absent from current sorters, absent from the analyses built on their output, and, arguably, absent from the general mindset of neural data analysis. What might help is well-calibrated posterior probabilities over the forms of ambiguity encountered along the pipeline, rather than assuming ambiguity only comes into play during clustering. One direction worth exploring is an end-to-end approach that learns waveform embeddings (Koukuntla et al., 2025) from ground-truth and simulated bursts, where posterior soft labels are explicitly modeled to be conservative around putative bursts and spike overlaps.

## Acknowledgments

We thank Asohan Amarasingham, Marcus Triplett, Charlie Windolf, Liam Paninski, and Nick Steinmetz for comments on earlier versions of this work. Daniel English was supported by NIH NS131858, NIH MH140076, the Whitehall Foundation, and the Simons Foundation. Sam McKenzie was supported by NIMH R01MH140076, the Whitehall Foundation, and R00MH118423-05S1.

## Supplementary Information

### Supplementary figures

Figure S1 illustrates the soft labeling procedure step by step (Supplementary Methods). Figure S2 shows juxtacellular data quality for each session. Figure S3 compares KS4 performance across ground-truth definitions, matching tolerances, and recording durations. Figure S4 splits KS4 performance by brain region. Figure S5 shows the robustness of KS4 performance to parameter tuning, extracellular SNR, and drift correction. Figure S6 shows the ISI distribution, ACG, and error-decomposed CCGs for every neuron. The sessions used are described in Table S1. For all sessions, the first and final few minutes of recording the juxtacellular unit were excluded, as the recording was particularly unstable.

**Table S1:** Recording statistics for each ground-truth neuron. Publicly released sessions are named by brain region: CA1, cornu ammonis 1; NEO, neocortex; SUB, subiculum; THA, thalamus. Duration, length of the analyzed recording. Reported values are median (95% interval across 1000 posterior resamplings of the ground-truth spike train). Burst fraction, fraction of spikes preceded by an interspike interval of at most 5 ms. Juxtacellular SNR, median spike height divided by the noise level (MAD/0.6745) of the 100-Hz high-passed juxtacellular trace. Extracellular SNR as in Fig. S5B. Putative cell type classification follows Petersen et al. (2021): interneuron (INT) if the extracellular trough-to-peak duration was at most 0.425 ms, or longer with an autocorrelogram rise time above 6 ms; pyramidal (PYR) otherwise. These criteria were used for cortex and hippocampus; thalamic neurons are not classified (–). (−). ^†^Uncertain: a high firing rate without bursts is atypical for a CA1 pyramidal cell.

| ID | Duration (min) | Rate (Hz) | Burst fraction | Juxta. SNR | Extra. SNR | Type |
|---|---|---|---|---|---|---|
| CA1-1 | 9.0 | 1.44 (1.42–1.46) | 0.191 (0.184–0.197) | 16.3 (16.2–16.4) | 5.36 (5.32–5.38) | PYR |
| CA1-2 | 10.2 | 5.29 (5.27–5.31) | 0.008 (0.008–0.010) | 15.9 (15.9–15.9) | 3.13 (3.11–3.16) | PYR |
| CA1-3 | 11.0 | 22.45 (22.34–22.55) | 0.010 (0.009–0.012) | 14.0 (14.0–14.0) | 6.44 (6.39–6.49) | INT |
| CA1-4 | 11.8 | 15.13 (15.11–15.14) | 0.000 (0.000–0.000) | 32.5 (32.5–32.5) | 2.36 (2.35–2.36) | PYR<sup>†</sup> |
| CA1-5 | 2.6 | 5.09 (5.00–5.18) | 0.339 (0.331–0.347) | 32.7 (32.6–33.0) | 5.36 (5.26–5.48) | PYR |
| CA1-6 | 14.4 | 5.86 (5.84–5.89) | 0.051 (0.049–0.054) | 35.7 (35.7–35.7) | 3.54 (3.51–3.57) | PYR |
| NEO-1 | 18.6 | 11.19 (11.15–11.23) | 0.056 (0.054–0.057) | 33.2 (33.1–33.2) | 7.09 (7.07–7.12) | PYR |
| NEO-2 | 5.8 | 9.00 (8.89–9.11) | 0.270 (0.266–0.274) | 19.6 (19.2–19.9) | 4.71 (4.65–4.78) | PYR |
| SUB-1 | 5.0 | 9.99 (9.93–10.04) | 0.323 (0.321–0.326) | 33.6 (33.6–33.7) | 4.10 (4.07–4.12) | PYR |
| SUB-2 | 15.2 | 3.54 (3.53–3.55) | 0.285 (0.284–0.286) | 23.8 (23.7–23.8) | 3.93 (3.93–3.94) | PYR |
| SUB-3 | 17.5 | 2.45 (2.44–2.45) | 0.000 (0.000–0.000) | 34.0 (34.0–34.0) | 3.33 (3.32–3.33) | PYR |
| THA-1 | 6.9 | 8.93 (8.91–8.95) | 0.045 (0.042–0.047) | 68.5 (68.4–68.5) | 4.80 (4.79–4.81) | – |
| THA-2 | 13.5 | 8.15 (8.11–8.19) | 0.146 (0.144–0.148) | 51.4 (51.3–51.4) | 4.07 (4.05–4.08) | – |
| THA-3 | 31.9 | 12.24 (12.23–12.25) | 0.095 (0.095–0.096) | 73.2 (73.2–73.3) | 3.01 (3.01–3.02) | – |
| THA-4 | 21.2 | 2.76 (2.75–2.77) | 0.041 (0.040–0.043) | 71.8 (71.8–71.8) | 19.55 (19.50–19.59) | – |
| THA-5 | 9.7 | 17.70 (17.66–17.73) | 0.042 (0.041–0.043) | 41.6 (41.6–41.6) | 3.54 (3.53–3.54) | – |

**Figure S1:**
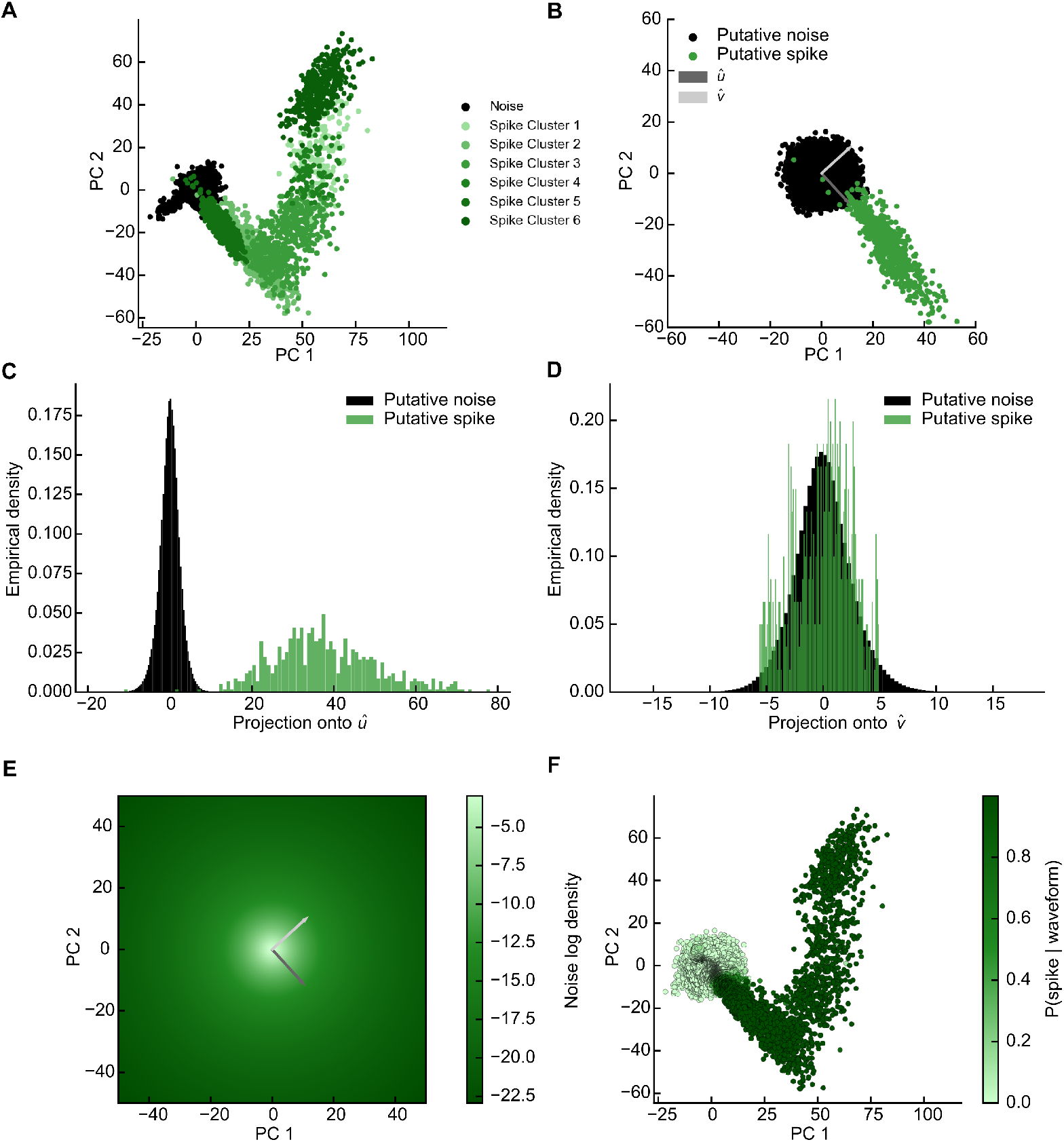
Technical details of soft assignments for juxtacellular recording. (A) After template learning and template matching, a BGMM is fit to putative noise and spike events in a 4-dimensional PCA basis (first two components shown); spike clusters are identified by criteria on their centroids. (B) Noise snippets are resampled across the recording (rejecting any snippets that overlap with a spike snippet from the BGMM fit). We assume the noise arises from an isotropic multivariate *t* distribution. A difference vector, ***û***, is computed between the noise mean vector and the putative spike centroid with the smallest norm. A vector ***v̂*** is then computed that (1) points in a non-spike-like direction (orthogonal to ***û***) and (2) maximizes the noise variance in the orthogonal subspace. (C) Along ***û*** the mode of the noise distribution is most decontaminated from spikes and we estimate its curvature. (D) Along ***v̂*** the tails of the noise distribution are most decontaminated from spikes, and we estimate the degrees of freedom *ν* by matching them to Student *t* tails. (E) Assuming isotropy, a multivariate *t* distribution is then implied in the *p*-dimensional PCA space. A class conditional density for spikes was already obtained from the BGMM. (F) Both class conditional densities are combined via Bayes’ rule, giving each event a posterior probability of being a spike, shown in the PCA space of (A) (see also Fig. 1C).

**Figure S2:**
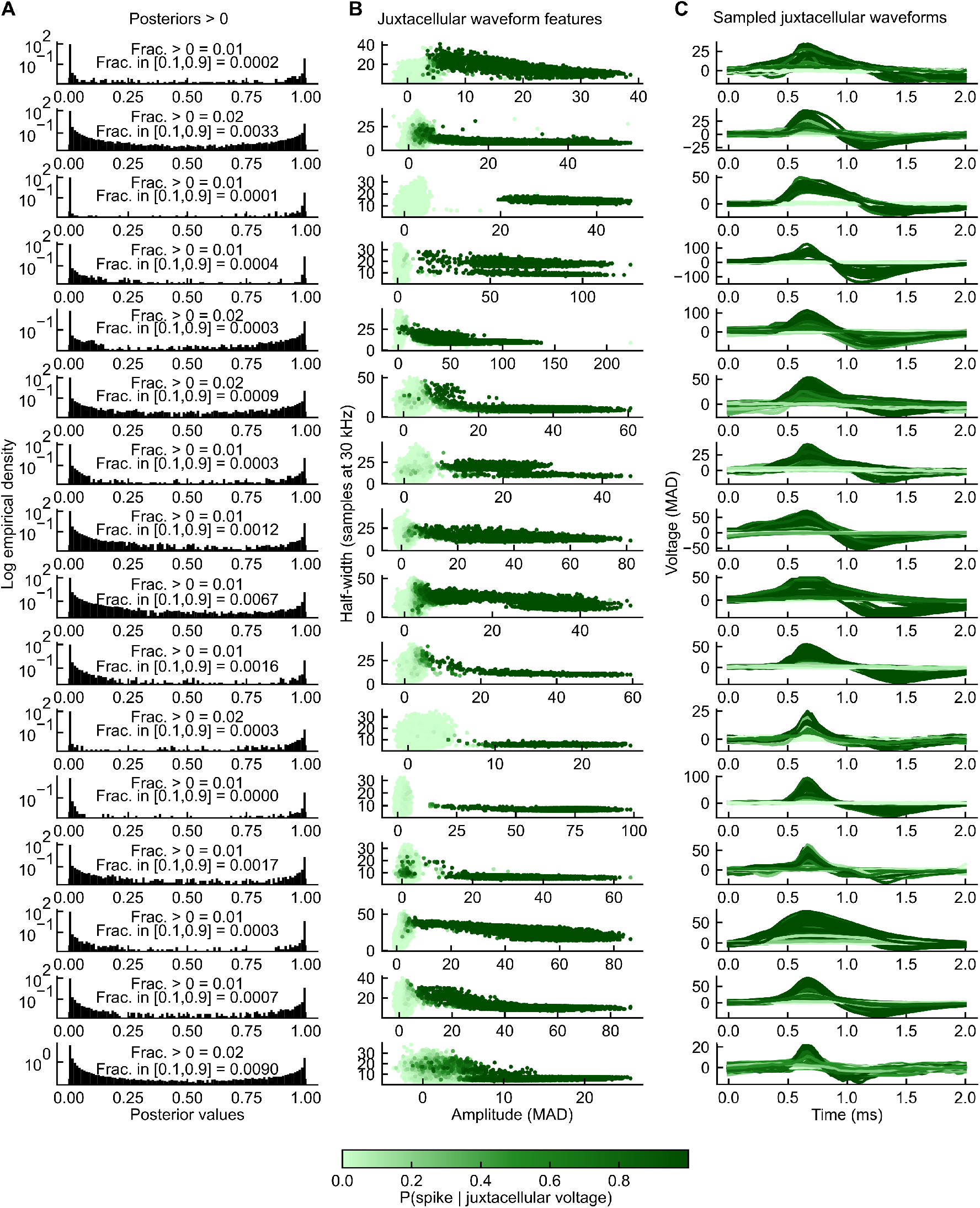
Juxtacellular data quality per session. One row per session (*n* = 16). (A) Distribution of posterior spike probabilities for candidate events with posterior *>* 0; note the log scale. Insets give the fraction of events with posterior *>* 0 and the fraction in the ambiguous range [0.1, 0.9]. (B) Half-width (samples at 30 kHz) versus amplitude (MAD-normalized) of juxtacellular events, colored by posterior probability of a spike. (C) Sampled juxtacellular waveforms, colored as in (B).

**Figure S3:**
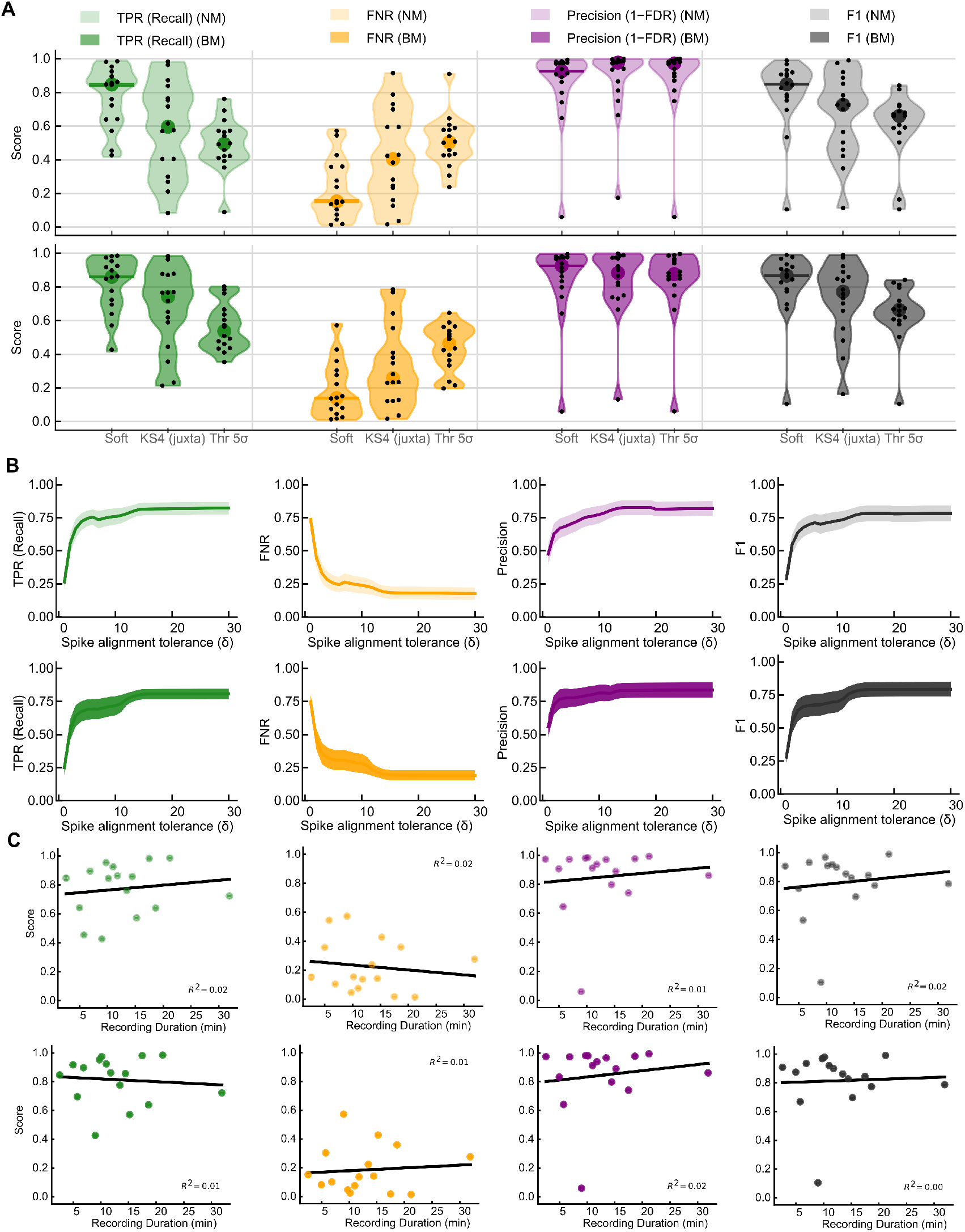
Sensitivity of KS4 performance estimates. (A) KS4 TPR, FNR, precision, and F1 evaluated against three ground-truth definitions: our soft labels, KS4 itself applied to the juxtacellular channel, and a 5*σ* amplitude threshold. Top, NM; bottom, BM; one dot per session. (B) Performance versus spike alignment tolerance *δ* (samples at 30 kHz); metrics saturate well before *δ* = 25 samples (≈0.83 ms), the value used throughout. Shading, SEM. Top, NM; bottom, BM. (C) Performance versus recording duration, one dot per session, with linear fits (all *R*^2^ ≤ 0.02). Top, NM; bottom, BM. All panels, *n* = 16 sessions.

**Figure S4:**
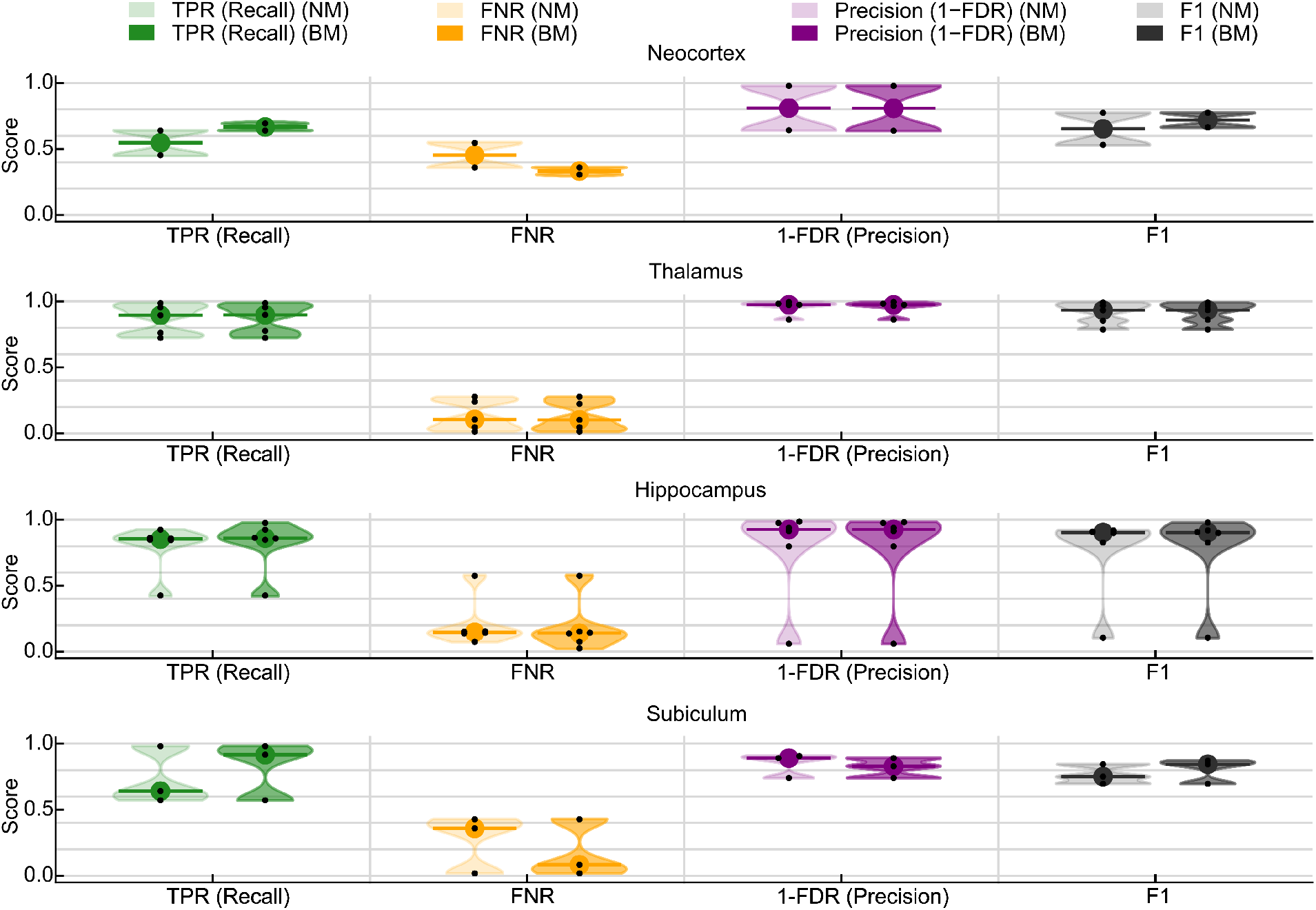
KS4 performance split by brain region. TPR, FNR, precision, and F1 for neo-cortex, thalamus, hippocampus (CA1), and subiculum, under NM (light) and BM (dark); dots, individual sessions; *n* = 16 sessions.

**Figure S5:**
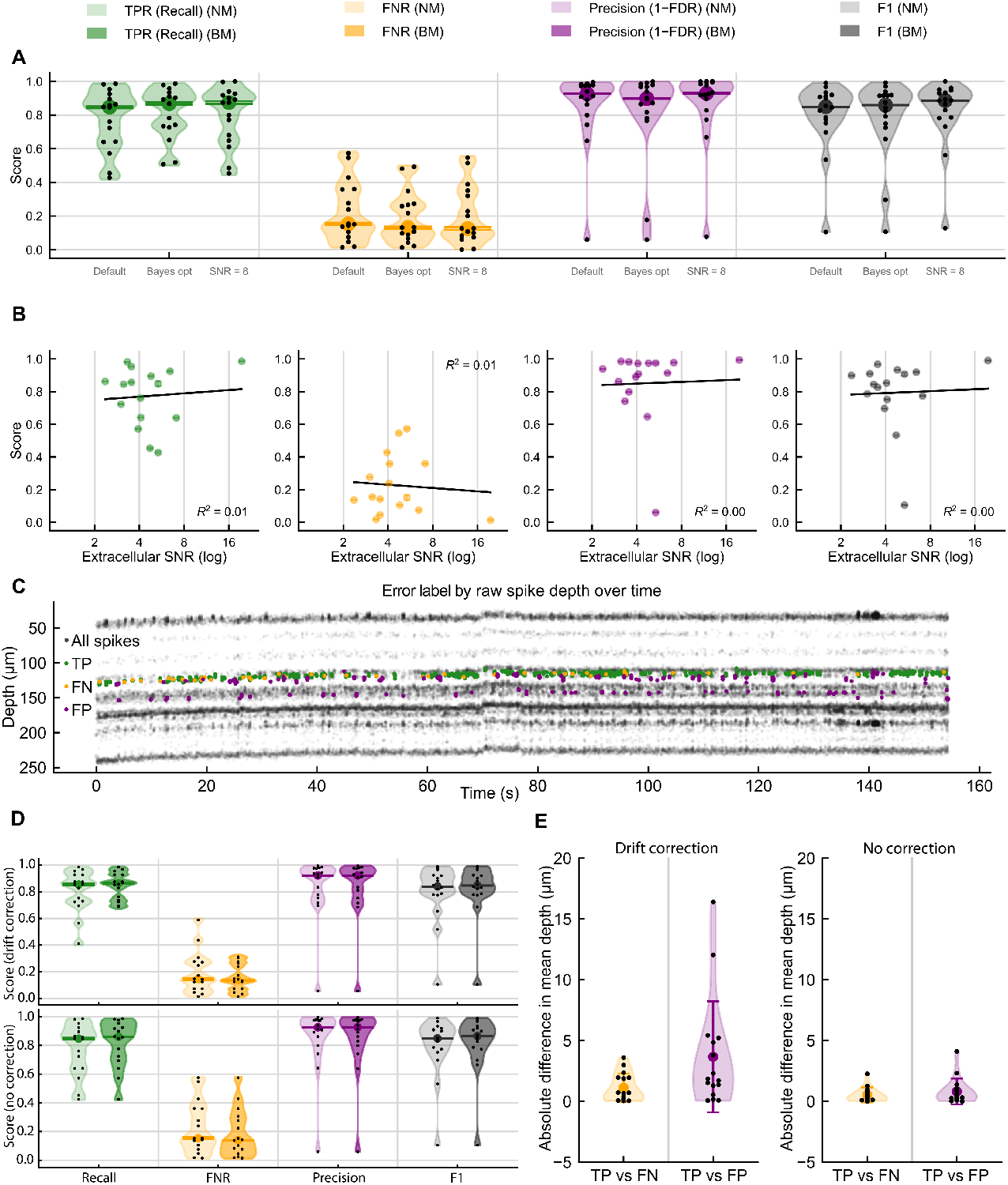
Robustness of KS4 performance evaluation to sorter parameters, extracellular SNR, and probe motion. (A) Overall performance of the KS4 NM unit with default parameters, with parameters tuned by Bayesian optimization (4-fold cross-validated; each session scored with parameters optimized on the other folds), and extrapolated to SNR = 8 (SpikeForest’s inclusion cutoff) by moving each neuron along a linear fit of the metric against SNR, refit for each surrogate. Violins, values pooled across surrogates; black dots, sessions; colored dot and bars, median and 95% interval of the across-session median across juxtacellular posterior surrogates. (B) Performance versus extracellular SNR (peak amplitude of the mean ground-truth waveform over noise on the peak channel, as in SpikeForest), one point per neuron; points, median across surrogates; bars, 95% interval; line, least-squares fit, with *R*^2^. (C) Example session showing sorting errors as the recorded neuron moved relative to the probe. Spike depth is raw, unregistered depth along the probe, so that motion remains visible; gray, all detected spikes, with TP, FN, and FP of the ground-truth unit overlaid. (D) Overall performance with (top) and without (bottom) drift correction, under NM (light) and BM (dark); one dot per session. (E) Spatial displacement of FN and FP from TP, from raw spike depths with drift correction enabled (left) or disabled (right); one dot per session; permutation test. Statistics in Supplementary Text; *n* = 16 sessions (one example session in C).

**Figure S6:**
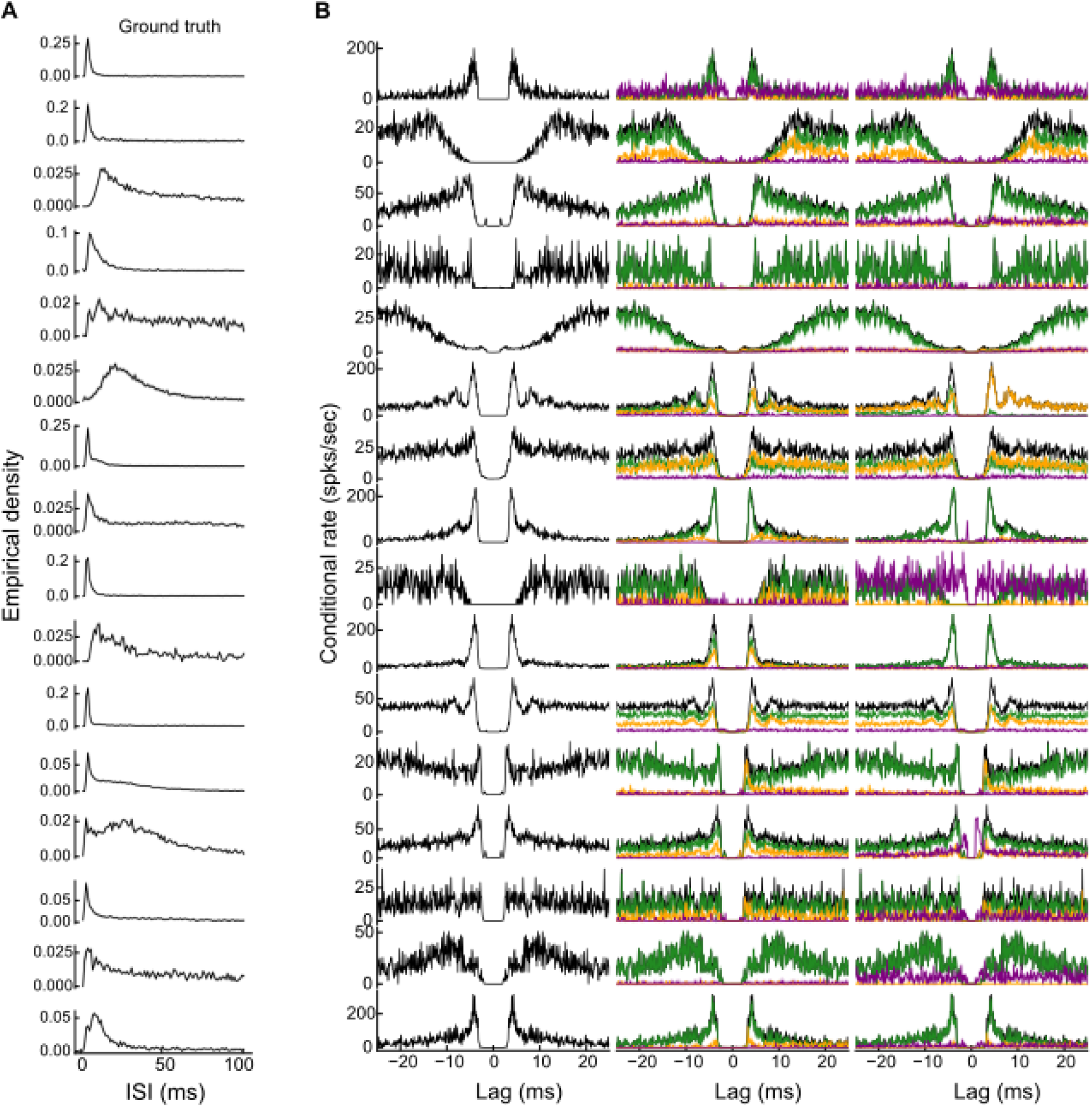
Spike train ISI distributions and ACGs (with error decompositions) for every ground-truth neuron. One row per neuron (*n* = 16). (A) Interspike interval (ISI) distribution of the ground-truth spike train. (B) Left, autocorrelogram (ACG) of the ground-truth spike train. Middle (NM) and right (BM), CCGs between the ground-truth spike train and its TP (green), FN (orange), and FP (purple) spikes, overlaid on the ACG (black).

## Supplementary Methods

### Soft labeling of juxtacellular spikes: full procedure

#### Preprocessing

The juxtacellular signal was high-pass filtered (third-order Butterworth, 100 Hz cutoff, zero-phase) and divided by a robust estimate of its noise standard deviation, *σ̂* = MAD */*0.6745, so amplitudes below are in MAD-normalized units.

#### Template learning

A canonical two-step procedure in spike sorting involves thresholding followed by feature extraction and clustering. Given our goal of modeling both spike and noise on a single channel, it is both feasible and desirable to reverse these steps: cluster first, then detect. Therefore, we rely on the intrinsic structure of the data to find clusters corresponding to spikes or noise and then identify spike-generating clusters based on their centroids.

Since spikes are rare events, the first problem is to correct for class imbalances. For each session, we set 100 equally spaced bins between zero and the maximum amplitude. Then, 100 events are sampled (with replacement) uniformly from each amplitude stratum. We z-score each sample of these snippets across events and project them into their 4-dimensional PCA basis, and perform an initial over-clustering with *k*-means (*k* = 8). PCA, *k*-means, Bayesian Gaussian mixture models and multi-task LASSO regression were all implemented with scikit-learn (Pedregosa et al., 2011).

Cluster centroids are accepted as spikes if (a) their amplitude at the alignment point exceeds 7 MAD-normalized units and (b) their peak lies within 5 samples of that point. Noise clusters are then discarded. We perform template matching with each accepted template and deduplicate detections in a winner-take-all fashion. In this pipeline, template matching is chiefly used to improve featurization by sharpening spike alignment.

#### Class conditional density for candidate spike events

A typical next step would be to find peaks in the template-matched signal and set a threshold for spikes. We found it more reliable and sensitive to instead recapitulate the stratified sampling step from before, but this time in terms of match strength rather than amplitude. For each session, we set 100 equally spaced bins between zero and the maximum match score. Then, 100 voltage snippets— sampled where the match score peaks—are sampled (with replacement) uniformly from each match score stratum.

We then perform Bayesian Gaussian mixture modeling (BGMM) in the 4-dimensional PCA basis of these data. Stratified sampling biases the training distribution from the full data distribution, and so a post-hoc correction is required. We assume the stratified sampling primarily biases mixture weights, not means and covariances.

The BGMM is fit with a Dirichlet process prior with a maximum of 8 components and a weight concentration prior favoring sparsity (*w*_0_ = 10^−5^). To correct for distributional shift, after training, we revise mixture weight estimates by elongating the EM algorithm as follows. For each of the *N*_test_ template-matched candidate events *i* (the full, unstratified set), we compute responsibilities *γ_i,m_* for each mixture component *m* and let 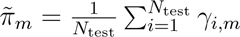. From these revised mixture weights, we obtain revised responsibilities *γ̃_i,m_*, and this update is iterated 10 times. Spike-generating clusters are identified based on their centroids as in the first pass.

At this point, the fitted Gaussian mixture covers a full range of noise and spike events, and we could compute the posterior probability that a data point belongs to any spike cluster. However, template matching is known to have poor selectivity: a high false positive rate is intrinsic to the procedure when Gaussian assumptions are violated, and particularly in high dimensions. Furthermore, because the base rate of noise is high (i.e., spikes are rare events) the precision (i.e., the fraction of correct positive detections) will be highly sensitive to the false positive rate. Since the goal here is to offer a ground-truth dataset, it makes sense to have high selectivity, and without making stronger assumptions about the neurons, this suggests a conservative noise model.

We therefore propose a post-hoc correction to the BGMM noise clusters. Identified noise or artifact clusters from the fitted BGMM are discarded, and the remaining mixture weights for the inferred spike clusters, indexed by *m* ∈ S, are renormalized to unity, giving weights *π_m_*; component *m* has mean ***µ****_m_* and covariance **Σ***_m_*. The remaining mixture now constitutes a class-conditional density for spike events.

#### Class conditional density for candidate noise events

We would now like to fit a robust model to the noise. We fit an isotropic multivariate *t* distribution using inference tactics similar in spirit to empirical Bayes. Using a multivariate *t* distribution has two motivations. Firstly, thinking of the juxtacellular recordings as “soft ground truth” to be employed for performance evaluation and model calibration for extracellular inference, it makes sense to have a conservative noise model. That is, all else being equal, the multivariate *t* distribution will place greater likelihood on noise at decision boundaries or outliers. Second, and more significantly, we observed in the data that random 2-dimensional projections of randomly sampled voltage snippets had isotropic heavy tails in sessions where the SNR was very large (i.e., when spike and noise clusters were highly separated).

Noise snippets were sampled at random times, excluding windows around detected spike events, and projected into the same PCA basis. In this *p*-dimensional basis (*p* = 4), let ***J****_t_* ∈ ℝ*^p^* denote the projected snippet at time *t*. We assume a noise event is distributed as ***J****_t_* | *noise* ∼ *t_p_*(***µ***_0_*, s*^2^***I****, ν*), an isotropic multivariate *t* distribution with location ***µ***_0_ (estimated as the median of the noise samples), scale *s*, and degrees of freedom *ν*. Its elliptical symmetry and closure under marginals imply that for any unit vector ***a*** ∈ ℝ*^p^*, ***a***^T^***J****_t_* ∼ *t*_1_(***a***^T^***µ***_0_*, s*^2^*, ν*): every 1-dimensional projection is a univariate *t* distribution with scale *s* and degrees of freedom *ν*.

Let ***µ****_m_*∗ be the mean of the spike component (from the previously fit BGMM) with the smallest norm, and let ***û*** = ***u****/*∥***u***∥, where ***u*** = ***µ****_m_*∗ − ***J̅***, with ***J̅*** the mean of the noise samples. We assume that if the noise samples are projected onto ***û*** the central region will be largely decontaminated from small spikes. Hence let *z* := ***û***^T^***J****_t_*, which has mode *µ_z_* = ***û***^T^***µ***_0_ and density

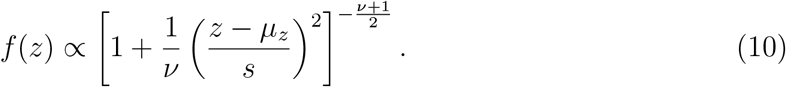

Taking the curvature of the log density at its mode and isolating *s*^2^ leads to the expression:

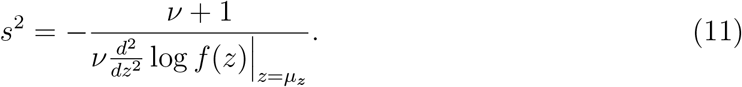

We estimate the curvature 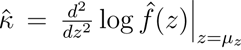 first through an estimate of *f̂*(*z*) via kernel density estimation (KDE) on samples of *z*, and then by evaluating the numerical second derivative around the inferred log density’s mode.

With *κ* estimated, the tails of the multivariate *t* distribution will then be affected by the value of *ν*, however the distribution of *z* is contaminated with spike events at its tails. Our goal is to estimate the tail shape of the null distribution without contamination from the signal cluster. Because signal and null differ primarily along ***û***, we expect signal contamination in the *null tails* to be minimal along directions orthogonal to this difference vector. To identify a projection direction that best decontaminates the null region from spikes and is a conservative estimate of spread, we seek a unit vector ***v̂*** that (1) lies orthogonal to ***û***, and (2) maximizes the variance of noise along that direction, thus minimizing contamination by spikes:

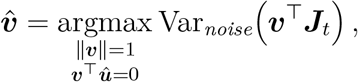

where the variance is taken over the noise samples. Let *P* = ***I*** − ***ûû***^T^ project onto the orthogonal complement of ***û***, and let ***C*** be the sample covariance of the noise samples. Then ***v̂*** is the eigenvector of *P* ***C****P* with the largest eigenvalue.

Analogously as before, let *y* = ***v̂***^T^***J****_t_*, and let *µ_y_* be its mean over the noise samples. For each candidate *ν*, the scale *s_ν_* is fixed by Eq. (11). Let *ĝ_ν_* be the empirical density of the standardized projection (*y* − *µ_y_*)*/s_ν_*, and let **Y** be its tails, the outer 5% of |*y* − *µ_y_*|*/s_ν_*. We choose

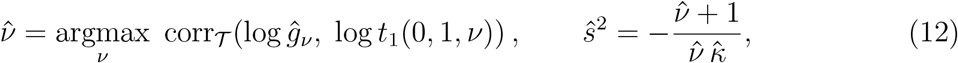

where *t*_1_(0, 1*, ν*) is the standard Student *t* density with *ν* degrees of freedom and corr**_Y_** is the Pearson correlation over the tails. The maximization is solved by grid search over *ν* ∈ [2, 50].

### Generative classifier

This use of a generative classifier is motivated by both the structure of the problem and theoretical work such as Ng and Jordan (2001). Ng and Jordan show that generative classifiers approach their asymptotic error with fewer training examples than discriminative ones, even though that asymptotic error is typically higher.

We are not trying to generalize to new recording sessions, but rather to describe uncertainty within a single dataset, where class boundaries may be ambiguous and expert-labeled data is scarce. A discriminative approach—for example, training a logistic regression or neural network to predict spike vs. noise based on our approximate labels—would commit to a fixed decision boundary optimized to match those labels. But in our setting, those labels are themselves only approximate, meaning a discriminative model may internalize label noise and reflect spurious confidence in regions where the boundary is ill-defined. Moreover, because such models are trained to optimize classification accuracy, not posterior fidelity, they may produce poorly calibrated probability estimates, especially near ambiguous cases where uncertainty matters most. In contrast, a generative model makes use of the full data distribution and allows us to incorporate domain-specific structure, such as assumptions about feature distributions or waveform variability, into the modeling process more transparently.

Under the hypotheses for a waveform ***J****_t_* at time *t*:

- *signal*: ***J****_t_* was generated by a spike from the recorded neuron.
- *noise*: ***J****_t_* was generated by background or unrelated processes.

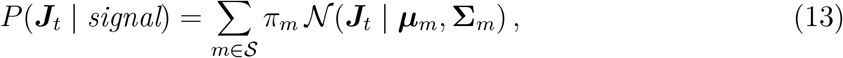

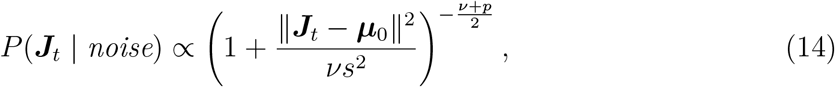

where *N*(· | ***µ***, **Σ**) is the multivariate normal density, *ν* and *s* take their estimated values, and the omitted normalizing constant, which depends on *s*, *ν*, and *p*, is included when computing the posterior below. Given a new voltage snippet at time *t*_0_, the posterior probability that ***J****_t_*_0_ is a spike is computed via Bayes’ rule:

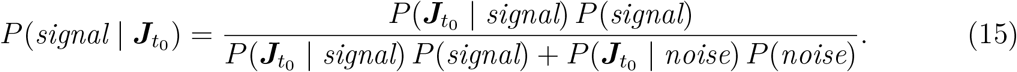

The prior *P* (*signal*) is set from the data as the fraction of the recording occupied by spike snippets: the number of candidate events assigned to a spike cluster, times the snippet length (60 samples), divided by the number of samples in the recording. *P* (*noise*) = 1 − *P* (*signal*).

#### Formal definitions of performance metrics

We now define performance evaluation metrics at the level of *extracellular* spike sorting, taking into account uncertainty at the juxtacellular level. Let *D* be a set of detected extracellular spike times postulated to correspond to the ground-truth neuron, and let *δ* be a time delay tolerance. A detected spike and a ground-truth spike match if they fall within *δ* of one another.

For surrogate *k*, recall is the fraction of sampled ground-truth spikes matched by some detected spike,

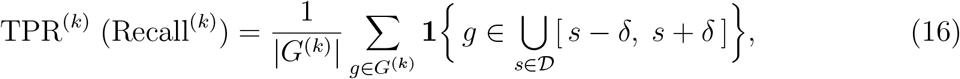

where the sum counts the TP spikes,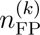, and 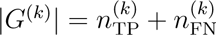;

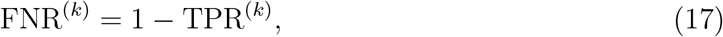

and the false discovery rate is the fraction of detected spikes not matched by any sampled ground-truth spike,

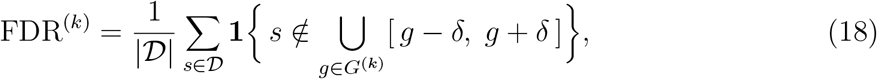

where the sum counts the FP spikes, 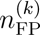.

For TPR and FNR, the normalization reflects how much juxtacellular spiking the Bayesian classifier believes exists. For FDR, the normalization reflects how much spiking the spike sorter purports; those claims are penalized in the numerator to the degree that detections are inconsistent with the juxtacellular model.

Precision and F1 follow for each surrogate *k*

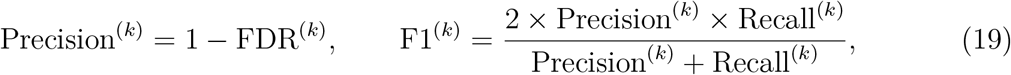

so all four metrics carry a distribution over the *K* sampled labelings rather than a single value.

#### Cross-correlogram definition

For two sets of event times *X* and *Y*, binned at Δ = 0.1 ms, the cross-correlogram (CCG) of *Y* relative to *X* is the conditional rate

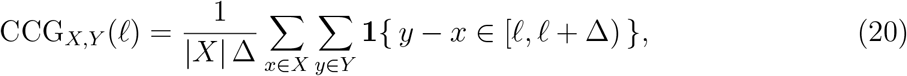

i.e. the rate of *Y* events at lag *ℓ* after an *X* event, for lags |*ℓ*| ≤ 25 ms. For surrogate *k*, the ACG takes *X* = *Y* = *G*^(^*^k^*^)^. For the TP, FN, and FP CCGs, *X* = *G*^(^*^k^*^)^ and *Y* is that surrogate’s set of TP, FN, or FP spike times, respectively; TP and FN times are ground-truth spike times, and FP times are the sorter’s detection times. Because TP and FN spikes are themselves members of *G*^(^*^k^*^)^, the zero-lag bin is excluded to remove each spike’s match with itself. CCGs were averaged across surrogates.

#### CCG asymmetry

Writing *C*(*ℓ*) for either CCG, the asymmetry is

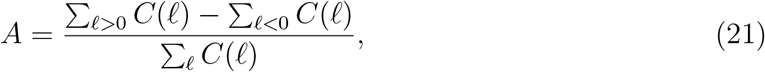

which is dimensionless and lies in [−1, 1]; a positive value indicates more of the CCG’s mass lies above zero lag.

#### Predictive comparison of intrinsic and extrinsic waveform variation

For each surrogate and sorter, let *y_i_* = 1 if ground-truth spike *i* was missed (FN) and *y_i_* = 0 if it was detected (TP). The two waveform predictors were the log-transformed intrinsic and extrinsic variation defined in the main Methods:

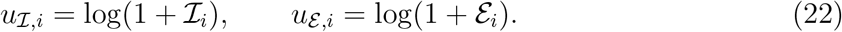

Within each classifier training fold, each included predictor was standardized using the mean and standard deviation estimated from that training fold. Denoting the resulting standardized predictors by *z_I,i_* and *z_ε,i_*, the four separately fitted models had linear predictors

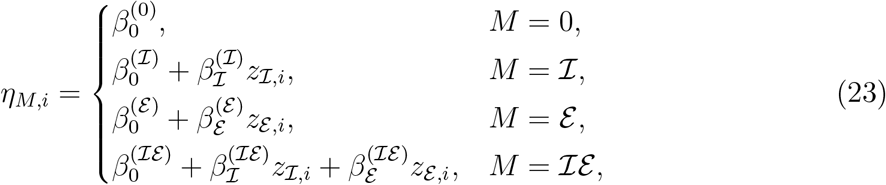

where *M* = 0, ***I***, *E*, and *IE* denote the intercept-only, intrinsic-only, extrinsic-only, and joint models, respectively. Under model *M*, the probability that event *i* was an FN was

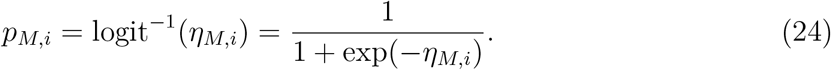

The three models containing waveform predictors were fit using L2-regularized logistic regression with inverse regularization strength *C* = 1. The intercept-only model assigned each held-out event the FN prevalence estimated from its training fold and contained no event-level waveform predictors.

All four models were evaluated using the same five stratified folds. Within each fold, the reference waveform **r**, predictor standardization, and logistic-regression coefficients were estimated from training-fold events only. Each event was therefore assigned a predicted FN probability by a model that was fit without that event.

Let *p̂_M,i_* denote the out-of-fold FN probability for event *i* from model *M* ∈ {0*, I, ε, Iε*}. For *N* sampled ground-truth events, the held-out log loss of model *M* was

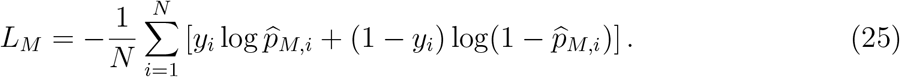

Lower log loss indicates more accurate probabilistic prediction of TP/FN status.

We quantified each predictor’s contribution to held-out predictive performance using two-feature Shapley contributions (Williamson and Feng, 2020). Because the reduction in log loss associated with one predictor can depend on whether the other predictor is included, its Shapley contribution was defined as the average reduction in held-out log loss across the two pairs of models that differed only in the inclusion of that predictor:

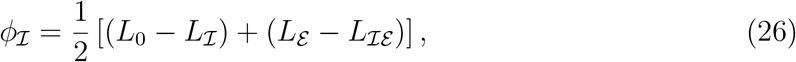

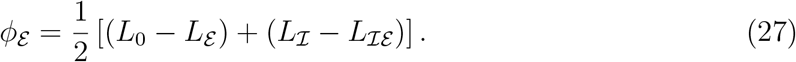

For *j* ∈ {*I, ε*}, a larger *ϕ_j_* indicates a greater contribution of predictor *j* to held-out prediction of TP/FN status. A positive *ϕ_j_* means that including predictor *j* reduced held-out log loss on average across the two relevant model comparisons; a negative *ϕ_j_* means that it increased held-out log loss on average. The two contributions sum exactly to the total held-out log-loss improvement of the joint model over the intercept-only model:

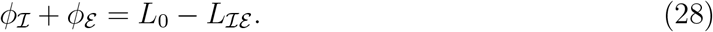

The logistic models and Shapley contributions were recomputed for each of the 1,000 surrogates without refitting the waveform LASSO. For each neuron, sorter, and surrogate *k*, we calculated

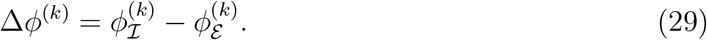

For each neuron and sorter, *ϕ**_I_***, *ϕ_ε_*, and Δ*ϕ* were summarized across surrogates by their medians; the 2.5th and 97.5th percentiles were additionally reported for the individual Shapley contributions. Across-neuron comparisons used each neuron’s median Δ*ϕ*. The group surrogate interval was obtained by calculating the median Δ*ϕ*^(^*^k^*^)^ across neurons separately for each surrogate and taking the 2.5th and 97.5th percentiles of those per-surrogate medians. Statistical significance was assessed using a one-sided exact paired sign-flip test on the per-neuron median differences, with Holm correction across the four sorters.

#### Exclusion criteria

The predictive comparison of intrinsic and extrinsic waveform variation (Fig. 6E,F) was restricted to neurons in which cross-fitted LASSO predictions reduced median held-out extracellular-waveform MSE relative to predicting the training-fold mean waveform; this criterion was based only on waveform prediction and was independent of the sorter-derived TP and FN labels. On the drift-off sort, 13 of 16 neurons met it; analyses based on distance and residual magnitude used all 16. Groups with fewer than 10 events were excluded from the matched-fraction comparisons of Fig. 5C. Substituting the mean waveform for a neuron failing the prediction criterion would effectively force intrinsic variation to zero while causing the residual to combine extrinsic variation, recording noise, and intrinsic variation not captured by the model; we therefore treat the intrinsic–extrinsic decomposition as unidentifiable for such neurons rather than as evidence that intrinsic variability is absent.

**Table S2:**
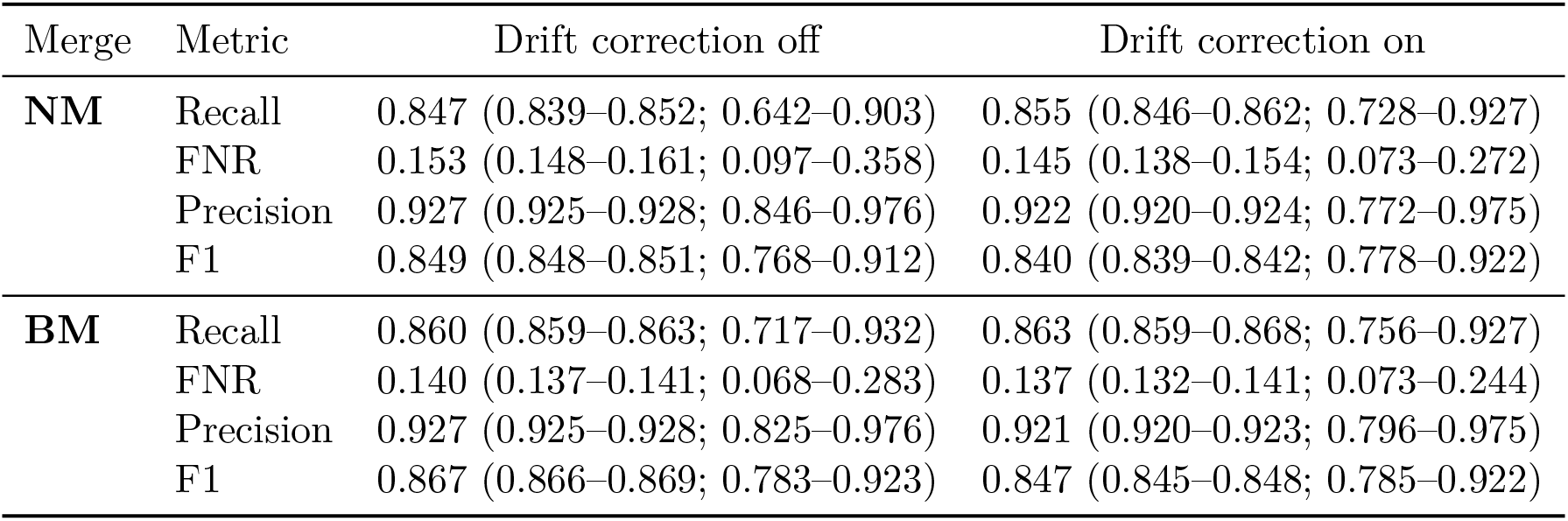
KS4 performance with and without drift correction. Values are reported as median (95% interval across surrogates; interquartile range across sessions). The drift-off column repeats Table 2.

| Merge | Metric | Drift correction off | Drift correction on |
|---|---|---|---|
| **NM** | Recall | 0.847 (0.839–0.852; 0.642–0.903) | 0.855 (0.846–0.862; 0.728–0.927) |
|  | FNR | 0.153 (0.148–0.161; 0.097–0.358) | 0.145 (0.138–0.154; 0.073–0.272) |
|  | Precision | 0.927 (0.925–0.928; 0.846–0.976) | 0.922 (0.920–0.924; 0.772–0.975) |
|  | F1 | 0.849 (0.848–0.851; 0.768–0.912) | 0.840 (0.839–0.842; 0.778–0.922) |
| **BM** | Recall | 0.860 (0.859–0.863; 0.717–0.932) | 0.863 (0.859–0.868; 0.756–0.927) |
|  | FNR | 0.140 (0.137–0.141; 0.068–0.283) | 0.137 (0.132–0.141; 0.073–0.244) |
|  | Precision | 0.927 (0.925–0.928; 0.825–0.976) | 0.921 (0.920–0.923; 0.796–0.975) |
|  | F1 | 0.867 (0.866–0.869; 0.783–0.923) | 0.847 (0.845–0.848; 0.785–0.922) |

## Supplementary text

### Sensitivity to ground-truth spikes rejected by the soft labeling

Because the soft labeling is deliberately conservative, some detections we count as FPs could be spikes of the ground-truth neuron that it rejected. If so, they would look like similar-template FPs: matched to the predicted waveforms. We tested this two ways. First, we asked how many FPs lie within *δ* of a candidate juxtacellular event that the labeler considered and rejected (posterior *>* 0.05): in the median neuron, 4% of its FPs, and no more among clean FPs than among coincident FPs. Second, we asked how many FPs lie within ±1 ms of any crossing of a permissive 5*σ* amplitude threshold on the juxtacellular trace (Quiroga et al., 2004), the most permissive labeling in Fig. S3A: in the median neuron, 6% of its FPs. These FPs matched the predicted waveforms no more often than the other FPs did (median fraction matched 0.37 vs. 0.30, in the 8 neurons with at least 10 of each; *p* = 0.27) and far less often than TPs (0.40 vs. 0.71, lower in 10 of 11 neurons, *p* = 0.003), whereas rejected ground-truth spikes should match about as often as TPs do; so most of them are ordinary FPs that happen to coincide with juxtacellular noise crossing the threshold. Removing them changed no conclusion: FPs remained coincident more often than TPs (86% vs. 70% of events in 13 of 15 neurons, against 83% vs. 70% with all FPs), and precision remained higher for clean than for coincident detections (0.97 vs. 0.92 in 13 of 15 neurons, against 0.96 vs. 0.92 in 12 of 15). We therefore report the analyses on all FPs.

### Probe and brain motion

KS4’s drift correction is not recommended for a probe of this geometry. We nonetheless enabled drift correction as a sensitivity check. Performance with drift correction enabled is compared with the default run in Table S2. With correction enabled, TPR rose slightly and precision fell slightly under both merge conditions, precision became more variable across sessions, and F1 was lower overall (Fig. S5D).

The spatial distribution of errors was consistent with this precision loss. To retain the physical motion visible in the recording, spike depth was analyzed in the raw, unregistered coordinate, obtained by adding KS4’s estimated drift back to its registered spike positions (Fig. S5C). For each session, we quantified spatial displacement as the absolute difference between the mean depth of true positives and the mean depth of false negatives or false positives. With drift correction enabled, false positives were displaced farther from true positives than false negatives by 2.54 *µ*m on average (two-sided permutation test, 10,000 permutations, *p* = 0.020; Fig. S5E). Without correction, the corresponding difference was only 0.28 *µ*m and was not significant (*p* = 0.421). Thus, drift correction modestly improved recall while admitting detections from more spatially displaced locations, degrading precision sufficiently to lower F1.

### Comparison with hard ground-truth definitions

To assess how the choice of ground-truth definition affects measured performance, we compared the soft labeling against two definitions that assign each detected event a hard label: KS4 applied to the juxtacellular channel (all units pooled) and a MAD-based amplitude detector at 5*σ* (Quiroga et al., 2004). For the single best-matched unit (NM), soft precision (median 0.927) was 0.010 lower than precision under the juxtacellular KS4 labeling (95% CI 0.004–0.018; 16/16 sessions, *p* = 3 × 10^−5^, Wilcoxon signed-rank) and 0.006 lower than under the 5*σ* detector (95% CI 0.004–0.011; 16/16 sessions, *p* = 3 × 10^−5^). For the best-merge condition (BM), soft precision was 0.004 lower than under the juxtacellular KS4 labeling (*p* = 0.82) and 0.001 lower than under the 5*σ* detector (*p* = 0.94), neither distinguishable from zero. Soft recall was higher than both alternatives. For NM, it exceeded recall under the juxtacellular KS4 labeling by 0.14 (*p* = 2 × 10^−4^) and under the 5*σ* detector by 0.32 (*p* = 3 × 10^−5^); for BM, the corresponding differences were 0.05 (*p* = 1 × 10^−3^) and 0.30 (*p* = 3 × 10^−5^).

Across both conditions and both alternative labelings, the precision cost of the soft labeling was at most 0.010, roughly 1% of its median precision, while recall differences ranged from 0.05 to 0.32. Restricting the ground truth to confidently labeled spikes therefore makes recall an upper bound on KS4’s performance, while costing it almost nothing in precision. So the conservative noise model does not disadvantage the sorter in either direction. This was appropriate given that our objective was to understand—rather than benchmark—spike sorting errors, which requires restricting analysis to trustworthy juxtacellular events.

### Performance across brain regions

With as few as two neurons per region, these comparisons are descriptive only (Fig. S4). Under NM, KS4 performed best in thalamus (median F1 0.93) and CA1 (0.90) and worst in neocortex (0.65) and subiculum (0.75). In every region the deficit came from recall rather than precision: neocortical and subicular TPR were 0.55 and 0.64, while precision was at least 0.81 everywhere. Best-merge recovered much of the missing recall in subiculum (TPR 0.64 to 0.92) and some in neocortex (0.55 to 0.67), but changed little in thalamus or CA1. This might indicate that oversplitting contributes to missed spikes in some regions.

### Time delay tolerance

Performance metrics are measured as a function of the time delay tolerance between true and estimated spike times. For each curve, we normalized by its terminal value and then identified the smallest tolerance, *δ*_0_, at which the magnitude of the curve exceeded 99% of the magnitude of the terminal value. Under NM, *δ*_0_ ranged from 0.03 to 0.77 ms (mean 0.23, s.d. 0.21 ms); under BM, the range was the same (mean 0.23, s.d. 0.23 ms). Because every curve had saturated by 0.77 ms, we adopted a slightly more lenient *δ* ≈ 0.83 ms (25 samples at 30 kHz) for all analyses.

### Sensitivity checks for re-sorting after artificial ground-truth neuron removal

The re-sorts contained fewer units than the original sorts (median 54, IQR 48–70, vs. 92, IQR 67–107; fewer in 15 of 16 sessions), so nearby-spike counts (Fig. 5F) were compared within each sort rather than across sorts. For the mislabeled fraction of FNs (Methods), subtracting the rate at which random times lost a detection changed the estimate by at most 0.05 in any neuron (median 0.75, IQR 0.53–0.90, versus 0.77 without the correction). Estimating the mislabeled fraction without splitting by attenuation gave a similar result (median 0.73), since the fraction of TPs losing a detection was nearly the same in attenuated and full-height spikes (0.91 and 0.94).

## References

Ida E. J. Aasebø, Mikkel E. Lepperød, Maria Stavrinou, Sandra Nøkkevangen, Gaute Einevoll, Torkel Hafting, and Marianne Fyhn. Temporal processing in the visual cor-tex of the awake and anesthetized rat. eNeuro, 4(4):ENEURO.0059–17.2017, 2017. doi: 10.1523/ENEURO.0059-17.2017.

Brian D. Allen, Caroline Moore-Kochlacs, Jacob G. Bernstein, Justin P. Kinney, Jörg Scholvin, Luís F. Seoane, Chris Chronopoulos, Charlie Lamantia, Suhasa B. Kodandara-maiah, Max Tegmark, and Edward S. Boyden. Automated in vivo patch-clamp evaluation of extracellular multielectrode array spike recording capability. Journal of Neurophysiology, 120(5):2182–2200, 2018. doi: 10.1152/jn.00650.2017.

Costas A. Anastassiou, Rodrigo Perin, György Buzsáki, Henry Markram, and Christof Koch. Cell type- and activity-dependent extracellular correlates of intracellular spiking. Journal of Neurophysiology, 114(1):608–623, 2015. doi: 10.1152/jn.00628.2014.

Pierre F. Apostolides, Aaron D. Milstein, Christine Grienberger, Katie C. Bittner, and Jeffrey C. Magee. Axonal filtering allows reliable output during dendritic plateau-driven complex spiking in CA1 neurons. Neuron, 89(4):770–783, 2016. doi: 10.1016/j.neuron.2015.12.040.

Alessio P. Buccino and Gaute T. Einevoll. MEArec: A fast and customizable testbench simulator for ground-truth extracellular spiking activity. Neuroinformatics, 19:185–204, 2021. doi: 10.1007/s12021-020-09467-7.

Alessio P. Buccino, Cole L. Hurwitz, Samuel Garcia, Jeremy Magland, Joshua H. Siegle, Roger Hurwitz, and Matthias H. Hennig. SpikeInterface, a unified framework for spike sorting. eLife, 9:e61834, 2020. doi: 10.7554/eLife.61834.

Jason E. Chung, Jeremy F. Magland, Alex H. Barnett, Vanessa M. Tolosa, Angela C. Tooker, Kye Y. Lee, Kedar G. Shah, Sarah H. Felix, Loren M. Frank, and Leslie F. Greengard. A fully automated approach to spike sorting. Neuron, 95(6):1381–1394, 2017. doi: 10.1016/j.neuron.2017.08.030.

Marlene R. Cohen and Adam Kohn. Measuring and interpreting neuronal correlations. Nature Neuroscience, 14(7):811–819, 2011. doi: 10.1038/nn.2842.

Alexander S. Ecker, Philipp Berens, R. James Cotton, Manivannan Subramaniyan, George H. Denfield, Cathryn R. Cadwell, Stelios M. Smirnakis, Matthias Bethge, and Andreas S. Tolias. State dependence of noise correlations in macaque primary visual cortex. Neuron, 82(1):235–248, 2014. doi: 10.1016/j.neuron.2014.02.006.

Daniel F. English, Sam McKenzie, Talfan Evans, Kanghwan Kim, Euisik Yoon, and György Buzsáki. Pyramidal cell-interneuron circuit architecture and dynamics in hippocampal networks. Neuron, 96(2):505–520, 2017. doi: 10.1016/j.neuron.2017.09.033.

Michale S. Fee, Partha P. Mitra, and David Kleinfeld. Variability of extracellular spike waveforms of cortical neurons. Journal of Neurophysiology, 76(6):3823–3833, 1996. doi: 10.1152/jn.1996.76.6.3823.

Benoît Frénay and Michel Verleysen. Classification in the presence of label noise: A survey. IEEE Transactions on Neural Networks and Learning Systems, 25(5):845–869, 2014. doi: 10.1109/TNNLS.2013.2292894.

George L. Gerstein and Wesley A. Clark. Simultaneous studies of firing patterns in several neurons. Science, 143(3612):1325–1327, 1964. doi: 10.1126/science.143.3612.1325.

Espen Hagen, Torbjørn V. Ness, Amir Khosrowshahi, Christina Sørensen, Marianne Fyhn, Torkel Hafting, Felix Franke, and Gaute T. Einevoll. ViSAPy: A Python tool for biophysics-based generation of virtual spiking activity for evaluation of spike-sorting algorithms. Journal of Neuroscience Methods, 245:182–204, 2015. doi: 10.1016/j.jneumeth.2015.01.029.

Kenneth D. Harris, Darrell A. Henze, Jozsef Csicsvari, Hajime Hirase, and György Buzsáki. Accuracy of tetrode spike separation as determined by simultaneous intracellular and extracellular measurements. Journal of Neurophysiology, 84(1):401–414, 2000. doi: 10.1152/jn.2000.84.1.401.

Darrell A. Henze, Zsolt Borhegyi, Jozsef Csicsvari, Akira Mamiya, Kenneth D. Harris, and György Buzsáki. Intracellular features predicted by extracellular recordings in the hippocampus in vivo. Journal of Neurophysiology, 84(1):390–400, 2000. doi: 10.1152/jn.2000.84.1.390.

Darrell A. Henze, Kenneth D. Harris, Zsolt Borhegyi, Jozsef Csicsvari, Akira Mamiya, Ha-jime Hirase, Anton Sirota, and György Buzsáki. Simultaneous intracellular and extracel-lular recordings from hippocampus region CA1 of anesthetized rats. CRCNS.org, 2009.

K. Jannis Hildebrandt, Maneesh Sahani, and Jennifer F. Linden. The impact of anesthetic state on spike-sorting success in the cortex: A comparison of ketamine and urethane anesthesia. Frontiers in Neural Circuits, 11:95, 2017. doi: 10.3389/fncir.2017.00095.

Roger M. Hoffer. The importance of “ground truth” data in remote sensing. NASA Technical Report NTRS 19730007768, Purdue University / NASA, 1972.

International Brain Laboratory, Kush Banga, Julius Benson, Jai Bhagat, Dan Biderman, Daniel Birman, Niccolò Bonacchi, et al. Reproducibility of in vivo electrophysiological measurements in mice. eLife, 13:RP100840, 2024. doi: 10.7554/eLife.100840.

Eugene M. Izhikevich, Niraj S. Desai, Elisabeth C. Walcott, and Frank C. Hoppensteadt. Bursts as a unit of neural information: Selective communication via resonance. Trends in Neurosciences, 26(3):161–167, 2003.

James J. Jun, Catalin Mitelut, Chongxi Lai, Sergey L. Gratiy, Costas A. Anastassiou, and Timothy D. Harris. Real-time spike sorting platform for high-density extracellular probes with ground-truth validation and drift correction. bioRxiv, page 101030, 2017. doi: 10. 1101/101030.

Sai Koukuntla, Tate DeWeese, Alexandra Cheng, Robyn Mildren, Aamna Lawrence, Austin R. Graves, Kathleen E. Cullen, Jennifer Colonell, Timothy D. Harris, and Adam S. Charles. SLAy-ing oversplitting errors in high-density electrophysiology spike sorting. bioRxiv, page 2025.06.20.660590, 2025. doi: 10.1101/2025.06.20.660590.

Jin Hyung Lee, David Carlson, Hooshmand Shokri Razaghi, Weichi Yao, Georges A Goetz, Espen Hagen, Eleanor Batty, EJ Chichilnisky, Gaute T Einevoll, and Liam Paninski. Yass: yet another spike sorter. Advances in neural information processing systems, 30, 2017.

Michael S. Lewicki. A review of methods for spike sorting: The detection and classification of neural action potentials. Network: Computation in Neural Systems, 9(4):R53–R78, 1998. doi: 10.1088/0954-898X_9_4_001.

Jeremy Magland, James J. Jun, Elizabeth Lovero, Alexander J. Morley, Cole Lincoln Hur-witz, Alessio Paolo Buccino, Samuel Garcia, and Alex H. Barnett. SpikeForest, repro-ducible web-facing ground-truth validation of automated neural spike sorters. eLife, 9: e55167, 2020. doi: 10.7554/eLife.55167.

André Marques-Smith, Joana P. Neto, Gonçalo Lopes, Joana Nogueira, Lorenza Calcaterra, João Frazão, Danbee Kim, Matthew G. Phillips, George Dimitriadis, and Adam R. Kampff. Recording from the same neuron with high-density CMOS probes and patch-clamp: A ground-truth dataset and an experiment in collaboration. bioRxiv, page 370080, 2018. doi: 10.1101/370080.

Juan Martinez, Carlos Pedreira, Matias J. Ison, and Rodrigo Quian Quiroga. Realistic simulation of extracellular recordings. Journal of Neuroscience Methods, 184(2):285–293, 2009. doi: 10.1016/j.jneumeth.2009.08.017.

Kevin P Murphy. Probabilistic machine learning: Advanced topics. MIT press, 2023.

Joana P. Neto, Gonçalo Lopes, João Frazão, Joana Nogueira, Pedro Lacerda, Pedro Baião, Arno Aarts, Alexandru Andrei, Silke Musa, Elvira Fortunato, Pedro Barquinha, and Adam R. Kampff. Validating silicon polytrodes with paired juxtacellular recordings: Method and dataset. Journal of Neurophysiology, 116(2):892–903, 2016. doi: 10.1152/jn.00103.2016.

Andrew Y. Ng and Michael I. Jordan. On discriminative vs. generative classifiers: A comparison of logistic regression and naive Bayes. In Advances in Neural Information Processing Systems 14, pages 841–848, 2001.

Marius Pachitariu, Nicholas A. Steinmetz, Shabnam Kadir, Matteo Carandini, and Kenneth D. Harris. Fast and accurate spike sorting of high-channel count probes with KiloSort. In Advances in Neural Information Processing Systems, volume 29, 2016.

Marius Pachitariu, Shashwat Sridhar, Jacob Pennington, and Carsen Stringer. Spike sorting with Kilosort4. Nature Methods, 21:914–921, 2024. doi: 10.1038/s41592-024-02232-7.

Alexandre Payeur, Jordan Guerguiev, Friedemann Zenke, Blake A. Richards, and Richard Naud. Burst-dependent synaptic plasticity can coordinate learning in hierarchical circuits. Nature Neuroscience, 24(7):1010–1019, 2021. doi: 10.1038/s41593-021-00857-x.

Antonio Pazienti and Sonja Grün. Robustness of the significance of spike synchrony with respect to sorting errors. Journal of Computational Neuroscience, 21(3):329–342, 2006. doi: 10.1007/s10827-006-8899-6.

Fabian Pedregosa, Gaël Varoquaux, Alexandre Gramfort, Vincent Michel, Bertrand Thirion, Olivier Grisel, et al. Scikit-learn: Machine learning in Python. Journal of Machine Learning Research, 12:2825–2830, 2011.

Peter C. Petersen, Joshua H. Siegle, Nicholas A. Steinmetz, Sara Mahallati, and György Buzsáki. CellExplorer: A framework for visualizing and characterizing single neurons. Neuron, 109(22):3594–3608, 2021. doi: 10.1016/j.neuron.2021.09.002.

Jonathan W. Pillow, Jonathon Shlens, E. J. Chichilnisky, and Eero P. Simoncelli. A model-based spike sorting algorithm for removing correlation artifacts in multi-neuron recordings. PLoS ONE, 8(5):e62123, 2013. doi: 10.1371/journal.pone.0062123.

R. Quian Quiroga, Zoltan Nadasdy, and Yoram Ben-Shaul. Unsupervised spike detection and sorting with wavelets and superparamagnetic clustering. Neural Computation, 16(8): 1661–1687, 2004. doi: 10.1162/089976604774201631.

Alex Rodriguez and Alessandro Laio. Clustering by fast search and find of density peaks. Science, 344(6191):1492–1496, 2014.

Cyrille Rossant, Shabnam N. Kadir, Dan F. M. Goodman, John Schulman, Maximilian L. D. Hunter, Aman B. Saleem, et al. Spike sorting for large, dense electrode arrays. Nature Neuroscience, 19(4):634–641, 2016. doi: 10.1038/nn.4268.

S. Murray Sherman. Tonic and burst firing: Dual modes of thalamocortical relay. Trends in Neurosciences, 24(2):122–126, 2001. doi: 10.1016/S0166-2236(00)01714-8.

Joshua H. Siegle, Peter Ledochowitsch, Xiaoxuan Jia, Daniel J. Millman, Gabriel K. Ocker, Shiella Caldejon, et al. Reconciling functional differences in populations of neurons recorded with two-photon imaging and electrophysiology. eLife, 10:e69068, 2021. doi: 10.7554/eLife.69068.

Giulia L. B. Spampinato, Elric Esposito, Pierre Yger, Jens Duebel, Serge Picaud, and Olivier Marre. Ground truth recordings for validation of spike sorting algorithms. Zenodo dataset, 2018.

Nicholas A. Steinmetz, Peter Zatka-Haas, Matteo Carandini, and Kenneth D. Harris. Distributed coding of choice, action and engagement across the mouse brain. Nature, 576: 266–273, 2019. doi: 10.1038/s41586-019-1787-x.

Mircea Steriade. Neocortical cell classes are flexible entities. Nature Reviews Neuroscience, 5(2):121–134, 2004. doi: 10.1038/nrn1325.

Mircea Steriade, Igor Timofeev, and François Grenier. Natural waking and sleep states: A view from inside neocortical neurons. Journal of Neurophysiology, 85(5):1969–1985, 2001. doi: 10.1152/jn.2001.85.5.1969.

Carsen Stringer, Marius Pachitariu, Nicholas Steinmetz, Charu Bai Reddy, Matteo Carandini, and Kenneth D. Harris. Spontaneous behaviors drive multidimensional, brainwide activity. Science, 364(6437):eaav7893, 2019. doi: 10.1126/science.aav7893.

Manik Varma and Debajyoti Ray. Learning the discriminative power-invariance trade-off. In Proceedings of the IEEE International Conference on Computer Vision (ICCV), 2007. doi: 10.1109/ICCV.2007.4408875.

Brian D. Williamson and Jean Feng. Efficient nonparametric statistical inference on population feature importance using Shapley values. In Proceedings of the 37th International Conference on Machine Learning, volume 119 of Proceedings of Machine Learning Research, pages 10282–10291. PMLR, 2020.

Iain H. Woodhouse. On ‘ground’ truth and why we should abandon the term. Journal of Applied Remote Sensing, 15(4):041501, 2021. doi: 10.1117/1.JRS.15.041501.

Jasper Wouters, Fabian Kloosterman, and Alexander Bertrand. SHYBRID: A graphical tool for generating hybrid ground-truth spiking data for evaluating spike sorting performance. Neuroinformatics, 19:141–158, 2021. doi: 10.1007/s12021-020-09474-8.

Pierre Yger, Giulia L. B. Spampinato, Elric Esposito, Baptiste Lefebvre, Stéphane Deny, Christophe Gardella, Marcel Stimberg, Florian Jetter, Günther Zeck, Serge Picaud, Jens Duebel, and Olivier Marre. A spike sorting toolbox for up to thousands of electrodes validated with ground truth recordings in vitro and in vivo. eLife, 7:e34518, 2018. doi: 10.7554/eLife.34518.

